# Time-lapse Image Super-resolution Neural Network with Reliable Confidence Quantification for Optical Microscopy

**DOI:** 10.1101/2024.05.04.592503

**Authors:** Chang Qiao, Shuran Liu, Yuwang Wang, Wencong Xu, Xiaohan Geng, Tao Jiang, Jingyu Zhang, Quan Meng, Hui Qiao, Dong Li, Qionghai Dai

## Abstract

Single image super-resolution (SISR) neural networks for optical microscopy have shown great capability to directly transform a low-resolution (LR) image into its super-resolution (SR) counterpart, enabling low-cost long-term live-cell SR imaging. However, when processing time-lapse data, current SISR models failed to exploit the important temporal dependencies between neighbor frames, often resulting in temporally inconsistent outputs. Besides, SISR models are subject to inference uncertainty that is hard to accurately quantify, therefore it is difficult to determine to what extend can we trust the inferred SR images. Here, we first build a large-scale, high-quality fluorescence microscopy dataset for the time-lapse image super-resolution (TISR) task, and conducted a comprehensive evaluation on two essential components of TISR neural networks, i.e., propagation and alignment. Second, we devised a deformable phase-space alignment (DPA) based TISR neural network (DPA-TISR), which adaptively enhances the cross-frame alignment in the phase domain and outperforms existing state-of-the-art SISR and TISR models. Third, we combined the Bayesian training scheme and Monte Carlo dropout with DPA-TISR, developing Bayesian DPA-TISR, and designed an expected calibration error (ECE)minimization framework to obtain a well-calibrated confidence map along with each output SR image, which reliably implicates potential inference errors. We demonstrate the unique characteristics of Bayesian DPA-TISR underlie the ultralong-term live-cell SR imaging capability with high spatial fidelity, superb temporal consistency, and accurate confidence quantification on a wide variety of bioprocesses.

## Introduction

In the past two decades, optical super-resolution microscopy (SRM) has become an essential tool for life science research. However, the increase in spatial resolution with any SRM is accompanied by the tradeoffs with other imaging metrics such as speed and duration^1, 2^. Recent advancements in deep neural networks have demonstrated remarkable capability to transform low-resolution (LR) images to their super-resolution (SR) counterparts, thereby enabling instant single image super-resolution (SISR) without any modifications on optical setups and realizing ultra long-term live-cell SR imaging^3-12^. Nevertheless, there are two major limitations in existing SISR neural networks when applied to time-lapse data. Firstly, when applied to process time-lapse images which is common in biological research, SISR models cannot capture the temporal dependencies between adjacent frames, thus yielding inferior SR fidelity and temporally inconsistent inferences^5^. Secondly, existing SISR models only generate monochromatic intensity images of biological structures without providing a quantitative and reliable evaluation on output confidence^13^, which makes it equivocal whether one could trust those outputs or not^14^. These two defects severely impede the wide applications of SR neural networks in routine biological imaging experiments.

To address the aforementioned challenges, we first employed our home-built multimodality structured illumination microscopy (SIM) system to acquire a large-scale high-quality dataset for the biological time-lapse image super-resolution (TISR) task, named BioTISR. The BioTISR dataset contains thousands of well-matched LR-SR time-lapse image stacks with three different input signal-to-noise ratios (SNR) and five biological specimens (Methods), allowing us to systematically evaluate the state-of-the-art TISR neural networks. Instead of assessing numerous TISR models individually, we investigated two most essential parts of the TISR network architecture, temporal information propagation and neighbor feature alignment, using a custom-designed general TISR framework (Methods). During the evaluation, we found that existing mainstream feature alignment mechanisms such as optical flow^15^ cannot always perform correct alignment speculatively due to the rapid motion of biological structures with global inconsistency and the intrinsic photon noise in fluorescent images (Supplementary Fig. 1). To this end, inspired by the frequency shifting property of Fourier transform, that is, the spatial shifting of structures equals to phase changes in Fourier domain, we devised a deformable phase-space alignment (DPA) mechanism that is able to adaptively learn large and tiny motions in the phase space at a sub-pixel precision. Furthermore, resorting to the strong feature alignment capability of DPA, we proposed the DPA-based time-lapse image super-resolution network (DPA-TISR) model and demonstrated that DPA-TISR outperforms other state-of-the-art TISR models in terms of both fidelity and temporal consistency.

To conquer the second issue, we incorporated Bayesian deep learning^16^ and Monte Carlo dropout approach^17^ with the DPA-TISR neural network, dubbed Bayesian DPA-TISR, to characterize the aleatoric uncertainty and epistemic uncertainty^16^, respectively. Then by calculating local integration of the mixed probability distribution function for each pixel (Methods), confidence maps that quantitatively evaluate the reliability of the output images could be generated. However, both our results and previous literature indicate that deep neural networks tend to be overconfident^18^, that is, the predicted confidence is higher than the real one. To cope with this issue, we developed an iterative finetuning framework to minimize the expected calibration error (ECE), where the ECE is defined as the weighted average of the absolute differences of inference accuracy and confidence, and can be reduced by more than 5-fold with our methods. We demonstrate that the DPA-TISR and Bayesian DPA-TISR enable low-cost SR live imaging with ultrahigh spatiotemporal resolution, extended duration, and reliable confidence evaluation by visualizing and analyzing various intracellular organelle interactions in live cells.

## Results

### Evaluation of essential components for TISR neural network models

TISR or video super-resolution (VSR) neural network models are designed to leverage temporal neighbor frames to assist the super-resolution of the current frame, thereby expected to achieve better performance than SISR models^19^ (Supplementary Note 1). Although TISR models have been widely explored in natural image SR to improve video definition, whether such models could be applied to super-resolve biological images, i.e., enhancing both sampling rate and optical resolution, has been poorly investigated. Here, we employed the TIRF/GI-SIM and nonlinear SIM^20^ modes of our home-built multimodality SIM system (Multi-SIM) to acquire an extensive TISR dataset of five different biological structures: clathrin-coated pits (CCPs), lysosomes (Lyso), outer mitochondrial membranes (Mito), microtubules (MTs), and F-actin filaments (Extended Data Fig. 1). For each type of specimen, we generally acquired over 50 sets of raw SIM images with 20 consecutive timepoints at 2-4 levels of excitation light intensity (Methods). Each set of raw SIM images was averaged out to a diffraction-limited wide-field (WF) image sequence and was used as the network input, while the raw SIM images acquired at the highest excitation level were reconstructed into SR-SIM images as the ground truth (GT) used in network training. In particular, the image acquisition configuration was modified into a special running order where each illumination pattern is applied 2-4 times at escalating excitation light intensity before changed to the next phase or orientation, so as to minimize the motion-induced difference between WF inputs and SR-SIM targets (Methods).

To effectively utilize the temporal continuity of time-lapse data, the state-of-the-art (SOTA) TISR neural networks consist of mainly two important components^21, 22^: temporal information propagation and neighbor feature alignment. We selected two popular types of propagation approaches: sliding window (Fig. 1a) and recurrent network (Fig. 1b), and three representative neighbor feature alignment mechanisms: explicit warping using optical flow^15^ (OF, Fig. 1c), and implicit alignment by non-local attention^23, 24^ (NA, Fig. 1d) or deformable convolution^21, 25, 26^ (DC, Fig. 1e), resulting in six combinations in total. For fair comparison, we custom-designed a general TISR network architecture composed of a feature extraction module, a propagation and alignment module, and a reconstruction module (Extended Data Fig. 2), and kept the architecture of the feature extraction module and reconstruction module unchanged while only modified the propagation and alignment module during evaluation (Methods). We then examined the six models on five different data types: linear SIM data of MTs, Lyso, and Mito, three of the most common biological structures in live-cell experiments; nonlinear SIM data of F-actin, which is of the highest structural complexity and upscaling factor in BioTISR; and simulated data of tubular structure with infallible GT references (Supplementary Note 2). As is shown in Fig. 1f, Extended Data Figs. 3 and 4, all models denoised and sharpened the input noisy wide-field (WF) image evidently, of which the model constructed with recurrent scheme and deformable convolution alignment resolved the finest details compared to the GT-SIM image (indicated by while arrows in Fig. 1f). Furthermore, we calculated time-lapse correlation matrixes (Fig. 1g) and image fidelity metrics (Fig. 1h-j), i.e., peak signal-to-noise ratio (PSNR) and structural similarity (SSIM), for the output SR images to quantitatively evaluated the temporal consistency and reconstruction fidelity, respectively. According to the evaluation, we found that (i) recurrent network-based propagation outperformed the sliding window-based one in both temporal consistency and image fidelity, and (ii) alignment mechanisms had little effect on temporal consistency of the reconstructed SR time-lapse data, while the DC-based alignment generally surpassed the other two mechanisms in terms of PSNR and SSIM for all types of datasets.

**Fig. 1.**
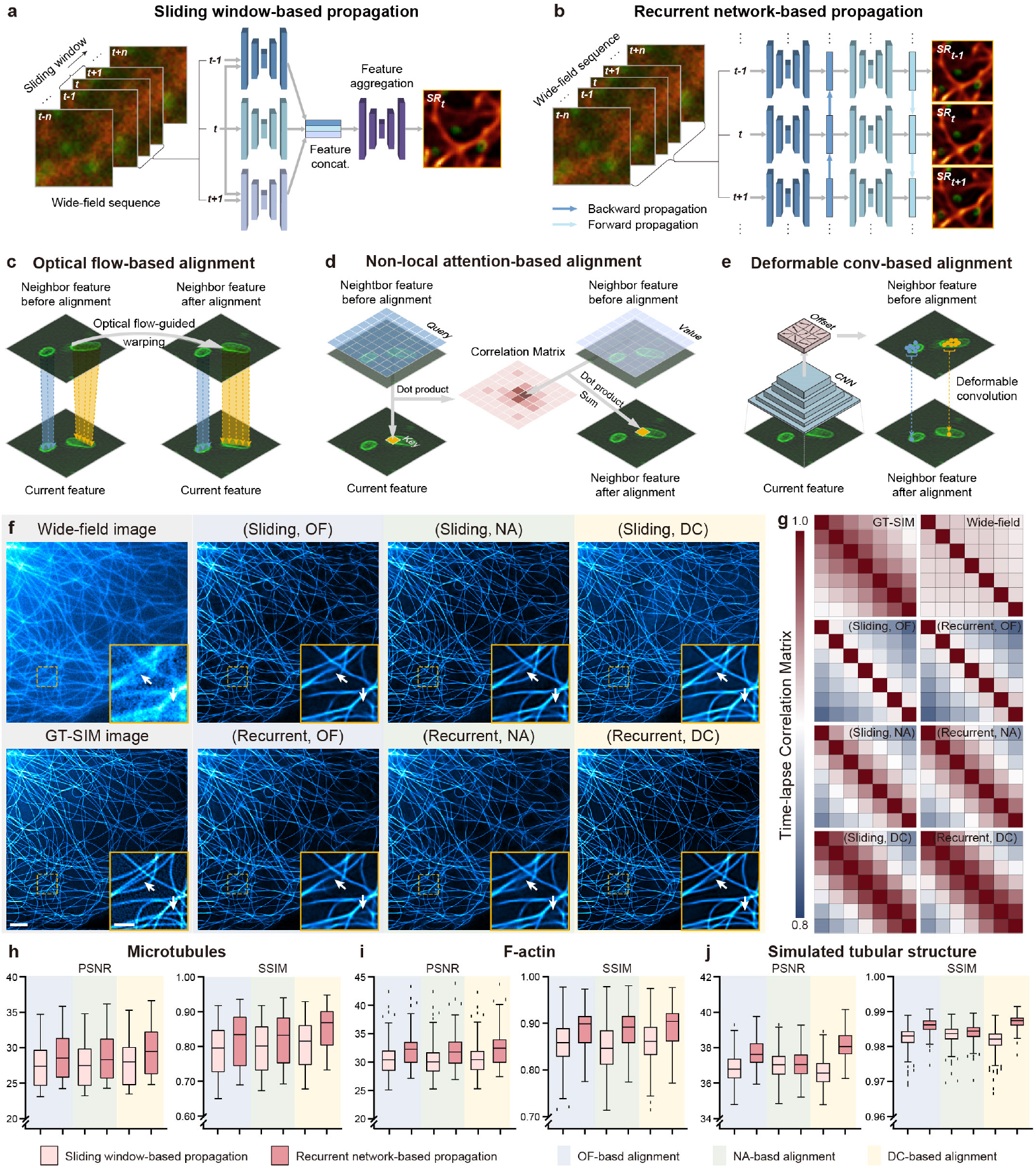
Comparison of representative propagation and alignment mechanisms in TISR models. **a-b**, Schematic illustration of two temporal information propagation mechanisms: sliding window-based propagation (Sliding, a) and recurrent network-based propagation (Recurrent, b). **c-e**, Schematic illustration of three neighbor feature alignment mechanisms using optical flow (OF, c), non-local attention (NA, d), and deformable convolution (DC, e). **f**, Representative TISR microtubule images inferred by six models combined by two propagation methods (Sliding and Recurrent) and three alignment mechanisms (OF, NA, and DC). WF and GT-SIM images are shown in the first column for reference. **g**, Time-lapse correlation matrixes of TISR images inferred by the models evaluated. **h-j**, Statistical comparison of the six models in terms of PSNR, SSIM on F-actin (h, n=50), microtubules (i, n=50), and simulated tubular structures (j, n=200). Scale bar, 3 μm (f), 1 μm (zoom-in regions in f).

Moreover, we validated these findings by using simulated dataset under different dynamic and noise conditions, i.e., structures with larger or smaller displacement between adjacent frames, and lower input SNR, with all experiments presented similar results (Supplementary Figs. 2-5). We speculate the underlying reasons are threefold: first, the bidirectional and recurrent propagation allows the TISR model to learn longer-range dependencies and maximizes the temporal information gathering compared to the sliding window with a limited size; Second, biological images contain heavier photon noises and more rapid changes between adjacent frames than natural images, which significantly interferes with the accuracy of OF calculation, leading to adverse impact for the OF-based alignment; Third, the NA-based alignment mainly mix-ups the information from spatially and temporally neighbor pixels, without modeling sub-pixel changes. In contrast, the DC-based alignment consists of both explicit sub-pixel shift estimation and implicit feature refinement, yielding the best capability to handle the complex motion pattern and spatially diverse speed for biological structures.

### Deformable phase-space alignment mechanism for time-lapse image super-resolution

Based on our comprehensive evaluation of the state-of-the-art methods, a strong baseline, which is a combination of recurrent network-based propagation and DC-based alignment, has been well-established. However, the existing DC mechanism only aggregate local features^26^, and fails to model global information and spatially consistent motion of biological structures. To this end, we further devised a deformable phase-space alignment (DPA) mechanism to enhance the global feature alignment at sub-pixel precision (Supplementary Note 3). In contrast to existing DC alignment which estimates offset in the spatial space, the proposed DPA primarily works in frequential space to adaptively learn phase residuals (Fig. 2a, Methods), which is corresponding to inflicting sub-pixel spatial shifting for each frequency component respectively. By visualizing the features before and after phase-space alignment, we showed that the phase-space alignment is able to fully exploit global dependencies to subtly model and compensate the movement of biological structures (Fig. 2b). Afterwards, we incorporated the superior feature alignment capability of DPA with the optimal TISR baseline model derived from the systematic evaluation to propose the DPA-based TISR neural network (DPA-TISR), as shown in Extended Data Fig. 5.

**Fig. 2.**
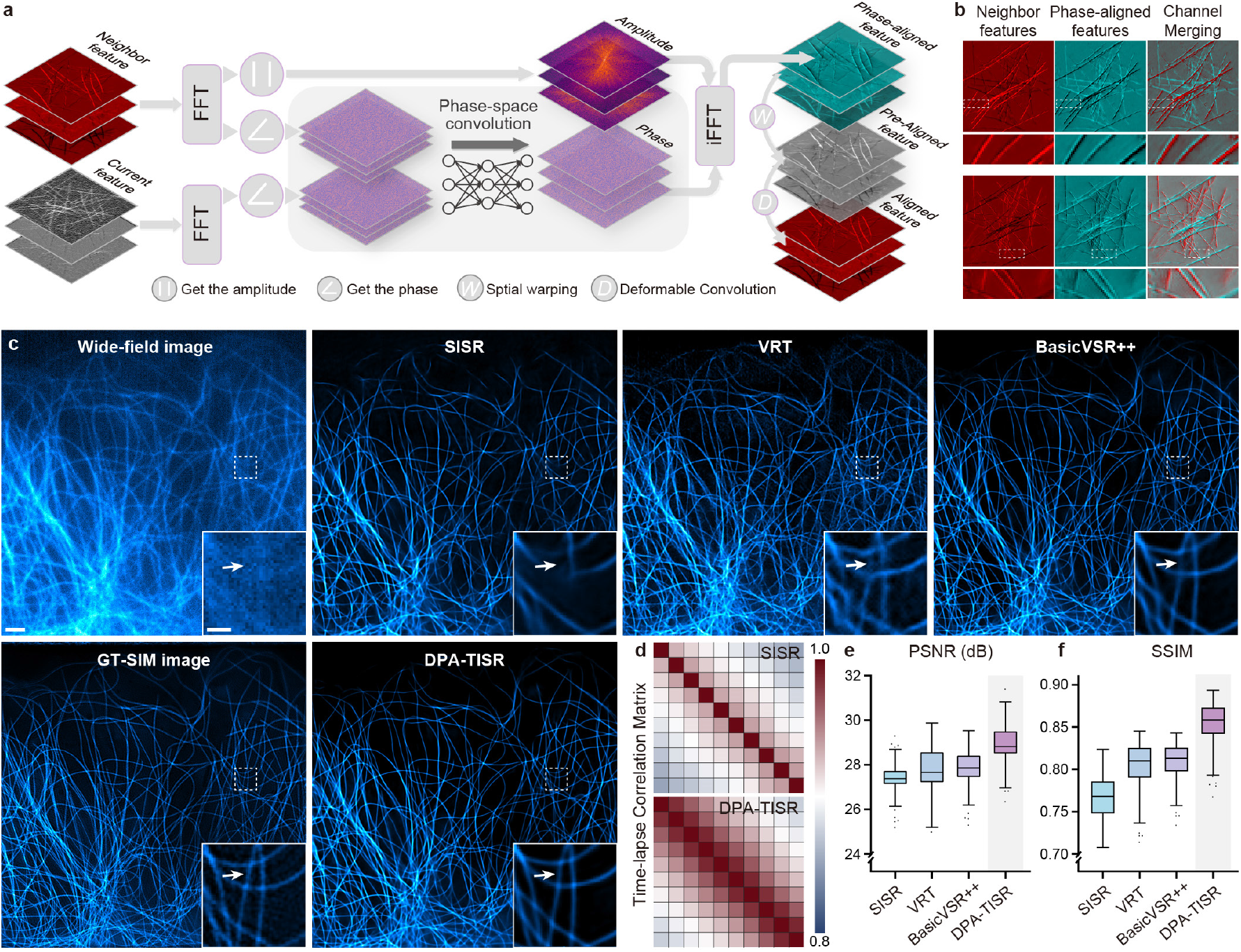
DPA mechanism and comparison of DPA-TISR with other methods. **a**, Schematic of the DPA mechanism. **b**, Visualization of representative features before and after phase-space alignment. **c**, Comparison of SR images of microtubules inferred by VRT, BasicVSR++, DPA-TISR, and a modified SISR model from DPA-TISR (Methods). WF and GT-SIM images are shown in the first column for reference. **d**, Correlation matrixes of SR time-lapse images inferred by the SISR model and DPA-TISR. **e**,**f**, Statistical comparison of VRT, BasicVSR++, DPA-TISR, and the modified SISR model in terms of PSNR (e) and SSIM (f), respectively (n=50). Scale bar, 3 μm (c), 1 μm (zoom-in regions in c).

To test whether DPA outperforms conventional spatial DA mechanism, we replaced the DPA by DA and two other variants of phase-space alignment, i.e., amplitude convolution and phase & amplitude convolution, in DPA-TISR (Supplementary Note 3). We found that the DPA with only phase convolution generally provided higher SR reconstruction fidelity in terms of PSNR and SSIM for both experimental and simulated dataset than other methods under similar computation complexity (Extended Data Fig. 6). Next, we compared the proposed DPA-TISR with two representative SOTA TISR models, i.e., BasicVSR++^21^, a superior baseline model combining the recurrent propagation and DC alignment, and VRT^27^, a recently published video restoration transformer that utilizes self-attention^28^ to model long-range temporal dependency, as well as a SISR variant of DPA-TISR (Methods) for biological time-lapse image super-resolution. As is shown in Fig. 2c-f, compared with TISR models, the SISR model failed to gathering cross-frame information thereby presented inferior noise robustness and output temporal consistency (Supplementary Fig. 6). Among three TISR models, with all aforementioned advances, DPA-TISR reconstructed finer details even in regions with severe noisy and background fluorescence than BasicVSR++ and VRT (Fig. 2c) and achieved the highest PSNR and SSIM (Fig. 2e, f).

### Rapid, long-term, SR visualization of organelle ultrastructure and dynamics by DPA-TISR

A great diversity of subcellular structures incessantly execute elaborate bioprocesses in living cells, among which F-actin cytoskeleton serves as a critical regulator of organelle positioning^29^ and is involved in various important cell functionalities such as clathrin-mediated endocytosis (CME)^30^. However, due to the extremely structural complexity of F-actin filaments, conventional SR live imaging methods have to impose relatively high excitation laser power to obtain enough fluorescence required by SR reconstruction, thus the multi-color SR observation of F-actin and other organelles is usually limited to ∼400 timepoints even assisted by SOTA deep learning-based methods^5^. To test whether DPA-TISR is competent in the TISR live-cell imaging task under relatively critical imaging conditions, we trained two independent DPA-TISR models for F-actin and CCPs, respectively, using corresponding datasets in BioTISR, and applied them to characterize the fast and subtle dynamics of F-actin filaments and CCPs from noisy time-lapse total internal reflective fluorescence (TIRF) images. As shown in Fig. 3a, although the noisy images were acquired using 10-fold lower excitation laser power than routine TIFR experiments, DPA-TISR successfully resolved densely interlaced F-actin filaments and ring structures of CCPs, yielding an extended SR observation window of 4,800 timepoints (Supplementary Video 1). Benefiting from the well-aligned temporal information, DPA-TISR substantially outperforms its SISR counterpart in both output image quality (Fig. 3b) and temporal consistency (Fig. 3c). Fig. 3d showcases that although the SISR model could also reconstruct the hollow structure of CCPs from a single WF image, their shapes varied unnaturally along with time supposedly due to the random noises in each frame. In contrast, CCPs inferred by DPA-TISR maintained temporally consistent with higher reliability, clearly characterizing the spatiotemporal regulation and interaction between F-actin and CCPs during CME processes (Fig. 3e).

**Fig. 3.**
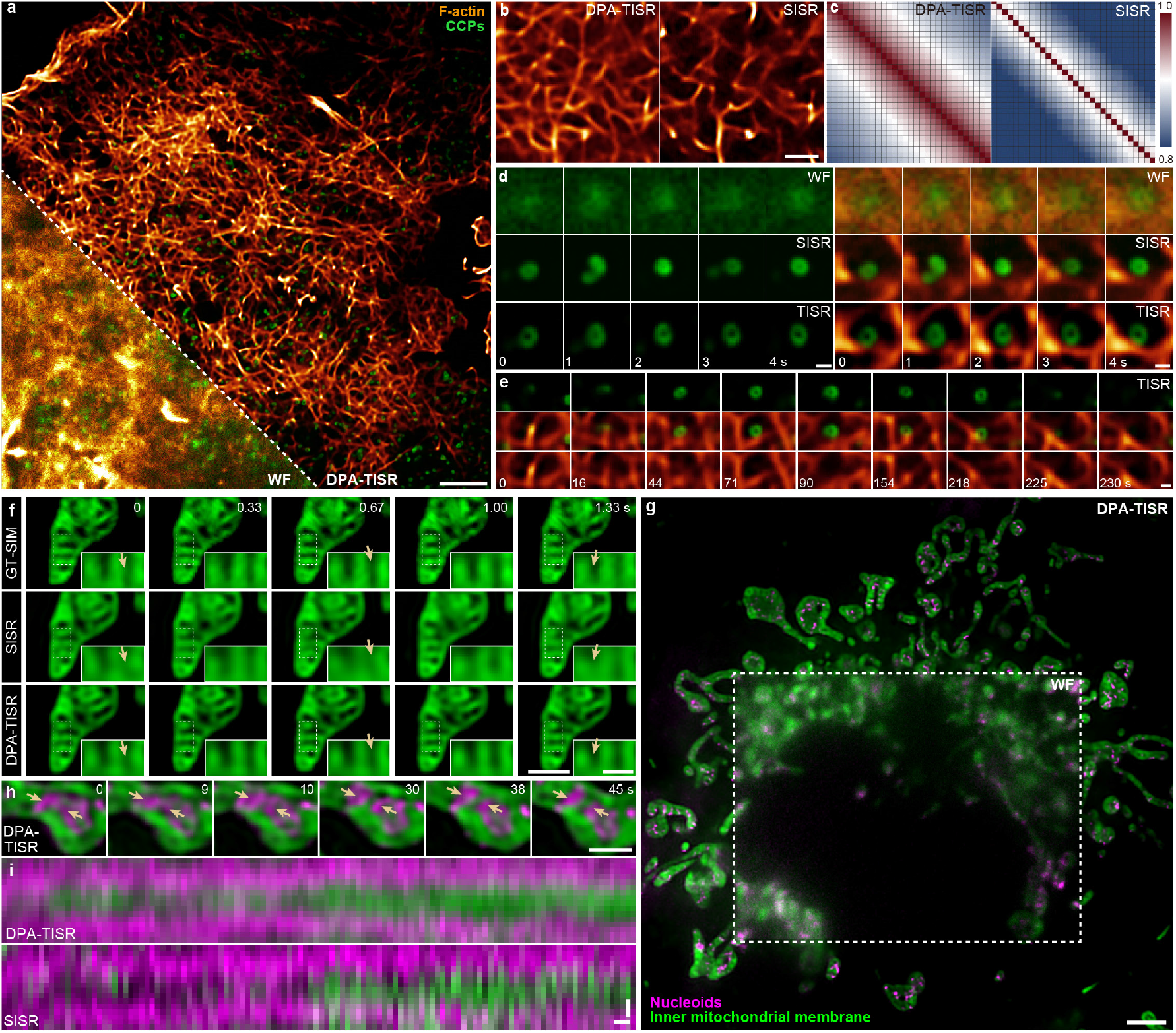
Rapid, long-term, SR visualization of organelle ultrastructure and dynamics by DPA-TISR. **a**, A representative SR frame of F-actin and CCPs reconstructed by DPA-TISR from a 4,800-frame two-color video (Supplementary Video 1). A fraction of wide-field (WF) image is displayed in the left bottom corner for comparison. **b**, Comparison of structural abundance of SR F-actin images inferred by the DPA-TISR model and its SISR counterpart (Methods). **c**, Correlation matrixes of SR time-lapse images inferred by the DPA-TISR and SISR models. **d**, Time-lapse SR images inferred by the DPA-TISR and SISR models showing the interactions between a CCP and F-actin filaments. **e**, Time-lapse DPA-TISR images showcasing the F-actin-CCPs contacts during clathrin-mediated endocytosis. **f**, Comparison of mitochondrial cristae time-lapse images inferred by SISR and DPA-TISR. The GT-SIM images are shown for reference. The yellow arrows indicate regions that are correctly inferred by DPA-TISR, whereas wrongly inferred by the SISR model. **g**, A representative SR frame of inner mitochondrial membrane and nucleoids reconstructed by DPA-TISR from a 1,680-timepoint two-color video (Supplementary Video 2). A fraction of WF image is displayed in the central region for comparison. **h**, Two-color time-lapse TISR images showcasing a typical nucleoid fission event concomitant with the formation of a new cristae in between. **i**, Kymographs drawn from SR images reconstructed via DPA-TISR and SISR models along the lines indicated by yellow arrowheads in h. Scale bar, 3 μm (a, g), 1 μm (b, f), 0.2 μm (d, e), 0.5 μm (zoom-in regions in f), 0.1 μm (horizontal bar in i), 1 s (vertical bar in i).

Besides the F-actin filaments, the mitochondrion is another exquisite organelle that exhibits a complicated inner architecture, i.e., the mitochondrial cristae with multiple nucleoids distributed in the interspace, of which the rapid dynamics are challenging to be long imaged using conventional SR microscopy^31, 32^. Although SISR neural networks can be utilized to reconstruct SR images from WF acquisitions, thus elongating the imaging duration, the overall time course has been limited to hundreds of timepoints^5^. To examine to what extend the DPA-TISR methodology could outperform the SISR scheme in this case, we individually trained two DPA-TISR models and two SISR models using dataset of inner mitochondrial membrane labelled with PKMO-Halo and nucleoids labelled by TFAM-mEmerald, respectively, then applied the well-trained models to process testing data (Fig. 3f) and a two-color time-lapse WF video of 1,680 timepoints that were acquired at low light conditions (Fig. 3g and Supplementary Video 2). We noticed that the cristae-like inner membrane invaginations inferred by the SISR model changed acutely among adjacent frames with lots of noise-induced errors, whereas the DPA-TISR model could take advantage of the temporal continuity of the time-lapse inputs and generated high-fidelity temporally-consistent SR results comparable to GT-SIM images (Fig. 3f). Facilitated by DPA-TISR, we clearly identified various delicate subcellular events such as nucleoid fission concomitant with the formation of a new cristae in between, even though they occurred in the maze-like ultrastructure of inner mitochondrial membrane (Fig. 3h and Supplementary Video 2).

### Confidence quantification and calibration with Bayesian DPA-TISR

Learning transformation from LR to SR images is essentially an ill-posed problem for either SISR or TISR. It means that there exist multiple solutions that correspond to the same LR inputs in the high-dimensional solution space^33^, and a neural network is just trained to extrapolate one statistically and perceptually good result with poor interpretability^34^. Therefore, in SR imaging experiments for scientific purpose, it is of vital importance to have an access to quantitatively evaluate the reliability of the network outputs^18, 35^. To this end, we incorporated Bayesian deep learning^16^ and Monte Carlo dropout^17^ with the proposed DPA-TISR, named Bayesian DPA-TISR, which is able to simultaneously reconstruct SR images and estimate corresponding uncertainty for each pixel of SR outputs (Fig. 4a, Methods, Supplementary Note 4). Generally, there are two major types of uncertainty that should be modelled^16^: the aleatoric uncertainty, also called data uncertainty, that arises from noise inherent in the observations; and the epistemic uncertainty, or called model uncertainty, which accounts for uncertainty in the model. Instead of directly inferring the fluorescence intensity values, Bayesian DPA-TISR is modified to estimate the Laplace distribution of each pixel, so as to capture the intrinsic aleatoric uncertainty (Methods). On the other hand, we introduce the dropout mechanism into the feature extraction module and reconstruction module of DPA-TISR, which allows Bayesian DPA-TISR to model the epistemic uncertainty and estimate the mixed probability distribution function (PDF) of each pixel by dropout-based self-ensemble (Methods). Moreover, parallel with the uncertainty characterization, a confidence map that intuitively indicates the output reliability could be generated by calculating the integral within an interval of the mixed PDF (Methods). The overall workflow of estimating uncertainty and generating confidence maps by Bayesian DPA-TISR is depicted in Fig. 4a.

**Fig. 4.**
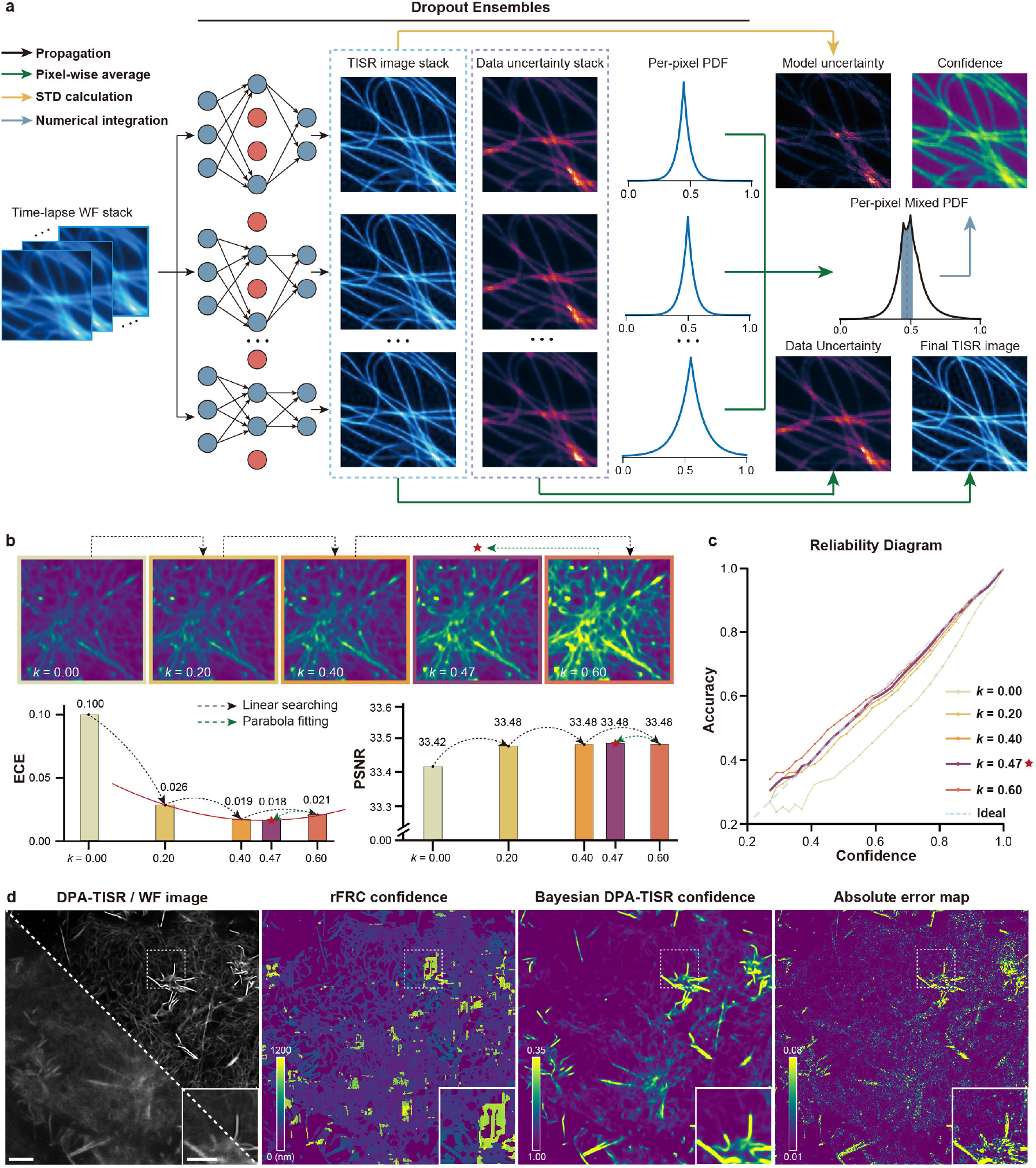
Confidence calculation and correction for DPA-TISR. **a**, Schematic illustration of uncertainty estimation and confidence map generation with Bayesian DPA-TISR. **b**, Progression of the confidence map (upper row), ECE value (lower left panel), and PSNR (lower right panel) during the iterative finetuning procedure of the Bayesian DPA-TISR. **c**, Reliability diagrams presented by accuracy curves versus average confidence for each searching step of iterative finetuning. **d**, Representative SR/WF image (first column) and confidence map generated by rFRC analysis (second column) and the well-calibrated Bayesian DPA-TISR model (third column). An absolute error map (fourth column) is shown for reference. Scale bar, 3 μm (d), 1 μm (zoom-in region in d).

However, after applying the Bayesian DPA-TISR to experimental data of multiple biological structures, we found that the estimated confidence is prone to be overconfident, i.e., higher than actual prediction accuracy (Extended Data Fig. 7), which is consistent with other literatures^18^. To conquer this issue, we devised an iterative finetuning framework to eliminate the expected calibration errors (ECE) between estimated confidence and inference accuracy. During the finetuning process, the objective function is defined as

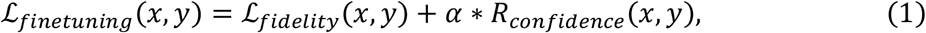

where *x* and y denote the TISR images inferred by Bayesian DPA-TISR and target GT-SIM images, respectively; ℒ_*fidelity*_ and ℒ_*confidence*_ are the fidelity loss and confidence correction regularization (CCR), and *α* is a weighting scalar to balance the two terms, which is empirically set as 0.1 in our experiments. The CCR is calculated by

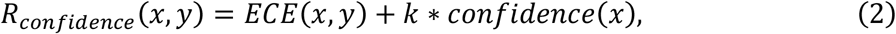

where *k* is the confidence suppression factor to be determined and optimized, and *confidence*(*x*) is the average confidence of the outputs. During the iterative finetuning procedure, the optimal value of *k* that minimizes the ECE will be determined by a combined strategy of linear searching and parabola fitting (Methods). As is shown in Fig. 4b, the optimal *k* can be determined with 3∼5 iterations, which typically takes less than one hour. With the proposed iterative finetuning process, the ECE can be reduced by more than five-fold, i.e., 0.1 to 0.018, while maintaining the output fidelity measured by PSNR, i.e., 33.48 dB v.s. 33.42 dB (Fig. 4b). By visualizing the reliability diagram that plots the accuracy, i.e., the fraction of successfully predicted pixels, versus average confidence, we found that the curve became substantially closer to the identity function after the iterative finetuning process (Fig. 4c). Fig. 4d shows the representative SR reconstruction, corresponding absolute error map, and confidence maps generated by Bayesian DPA-TISR after correction and rolling Fourier ring correlation (rFRC)^36^. It can be noticed that the predicted confidence map is highly consistent with the error map, while the latest rFRC methods gave a coarse confidence evaluation that is inconsistent with the actual error (Supplementary Note 5). Moreover, using datasets of multiple organelles, i.e., lysosomes, mitochondria, and F-actin in BioTISR, we validated that the proposed confidence correction framework could generally reduce ECE by 5∼10 fold with high generalization capability (Extended Data Figs. 7-9), implying that the calibrated confidence map estimated Bayesian DPA-TISR is well-competent to be a qualified error indicator.

### Long-term confidence-quantifiable SR live imaging via Bayesian DPA-TISR

Mitochondria are highly dynamic and undergo fission and fusion to maintain a functional mitochondrial network, of which the interactions with other organelles are essential for cellular homeostasis^37^ and quality control^38^. However, most literatures relied on conventional SR techniques to study these bioprocesses, e.g., interactions between mitochondria and lysosomes, with a narrow observation window of tens to hundreds of timepoints, which is limited by the phototoxicity and rapid photobleaching^37, 39^. To investigate the interaction dynamics over an extended time course, we acquired TIRF images using relatively low illumination intensity, and then reconstructed them into SR time-lapse images via well-trained DPA-TISR models. As such, we were able to record the interactions between Mito and Lyso in live COS-7 cells at a high spatiotemporal resolution for more than 10,000 two-color timepoints (Extended Data Fig. 10a and Supplementary Video 3), which is of two orders of magnitude more than conventional SIM imaging^37, 40^. Thus, it allowed for continuous capture of Lyso-Mito contacts and Lyso-mediated mitochondrial dynamics with quantitative confidence indicating the reliability of the events observed (Extended Data Fig. 10b-d and Supplementary Fig. 7).

Besides Lyso, the peroxisome (PO) is another organelle that has been reported to occasionally contact with Mito to exert its functionality in regulating overall cellular lipid and reactive oxygen species metabolism within mammalian cells^41^. However, the category and proportion of Mito-PO contacts have been rarely explored which is perhaps limited by live-imaging duration of existing SR techniques. To characterize the Mito-PO contacts, we integrated Bayesian DPA-TISR with our Multi-SIM system to record a COS-7 cell line labelled with 2xmEmerald-Tomm20 and PMP-Halo (Methods). Facilitated by the high-speed and long-term SR imaging capabilities, we were able to clearly distinguish the correlated interactions from random motions of the two organelles (Fig. 5a and Supplementary Video 4). In contrast, WF and conventional GI-SIM images with the same fluorescence budge are subject to diffraction-limited resolution and heavy noise-induced reconstruction artifacts, respectively (Fig. 5b). We tracked the displacements of individual POs and then calculated the Mander’s overlap coefficient (MOC) of each PO with its neighboring outer mitochondrial membrane for each frame to quantify the Mito-PO contacts (Methods). We found that nearly half of the identified POs (n=118 out of 210 from 3 cells) underwent no Mito-PO contacts (Fig. 5c), while the other half were closely associated with Mito, which could be further classified into three categories (Supplementary Video 5): 17% of POs (n=36) underwent stable association with contact sites from a single mitochondrion (Fig. 5d); 8% (n=17) of POs served as the bridge that simultaneously tethered two or more mitochondria (Fig. 5e); and 11% of POs (n=24) unexpectedly changed their contact sites from one mitochondrion to another, acting like intracellular messengers (Fig. 5f). In particular, we retained the last 7% POs (n=15) as unidentified (Fig. 5g) in that the Bayesian DPA-TISR alerted us that the output confidence of the underlying regions in Mito TISR images is too low to make accurate classifications (Fig. 5h). In brief, the Bayesian DPA-TISR enable the rational study of intricacy of complex Mito-PO contact behaviors. These observations demonstrate that Bayesian DPA-TISR lays the groundwork for opening up a wider application range of ultralong-term live-cell SR imaging and confidence-quantifiable biological analysis.

**Fig. 5.**
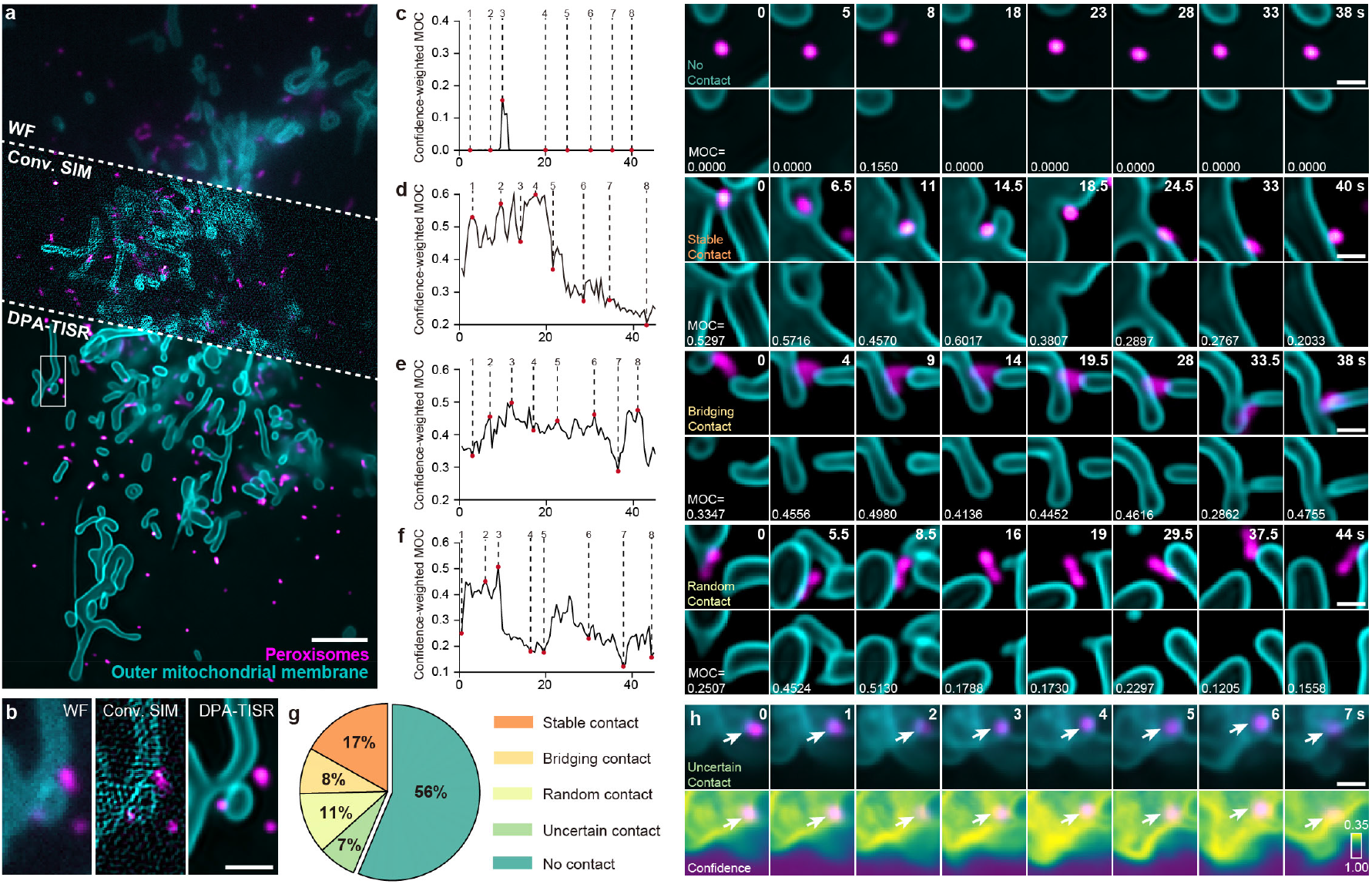
Long-term SR live imaging with reliable confidence evaluation via Bayesian DPA-TISR. **a**,**b**, Comparison of WF, Conv. SIM, and DPA-TISR images of outer mitochondrial membrane and peroxisomes. **c**-**f**, Dynamic characteristics of Mander’s over coefficient (MOC) for POs and corresponding time-lapse two-color TISR images of Mito and POs, showcasing four typical modes of Mito-PO contacts (Supplementary Video 5). **g**, Sector diagram of the proportion of POs in terms of their interactions with Mito. **h**, Time-lapse DPA-TISR images of Mito and POs (upper row) and corresponding confidence maps of Mito (lower row) with POs labelled with white arrows. Scale bar, 3 μm (a), 1 μm (b), 0.2 μm (c-f and h).

## Discussion

In this work, we first provided an open-access time-lapse SR image dataset named BioTISR, which is used to evaluate existing TISR neural network models and develop the DPA-TISR. We regard the BioTISR as a complementary extension of our previously released BioSR dataset^5^ and expect they will inspire and support more extensive developments of computational SR methods in the future. Next, we decoupled the TISR neural networks into two essential components, temporal information propagation and neighbor feature alignment, and systematically evaluated five representative mechanisms. With this comprehensive study, two optimal solutions for the above two processes were concluded, forming a strong baseline. Moreover, we devised the phase-space alignment mechanism as a complementary refinement in frequency domain for the spatial deformable convolution, and developed the DPA-TISR neural network model. We demonstrated that DPA-TISR could efficiently aggregate information of biological specimens from neighbor frames and generate time-lapse SR reconstructions of much superior fidelity and time-consistency compared with existing methods.

Furthermore, we upgraded DPA-TISR to Bayesian DPA-TISR that could generate a confidence map accompanied with its SR outputs. To repress the intrinsic over-confidence effect, we devised a combined loss consisting of an image fidelity term and a confidence calibration regularization term (Eqs. 1 and 2), as well as an iterative finetuning framework to minimize the expected errors between average confidence and inference accuracy. The finetuning procedure can be finished within 3∼5 steps with each step taking about 10 minutes. After optimization, the ECE of Bayesian DPA-TISR can be substantially reduced, resulting in a well-calibrated confidence quantification capability. With reliable confidence evaluation, Bayesian DPA-TISR enables error-aware SR investigation and analysis of intracellular interactions between mitochondria, cytoskeleton, and other organelles at an extended observation window up to 10,000 timepoints. These results demonstrate the potential of Bayesian DPA-TISR to greatly advance the application of SR microscopy in live-cell imaging.

Several improvements and extensions of DPA-TISR can be envisioned. It is known that deep neural networks are subject to the spectral bias towards low-frequency patterns^42^, which accounts for the resolution degradation of the output SR images. Adding regularization terms such as the discriminative loss^43^, perceptual loss^44^, and Fourier space loss^45^ during training procedures may be helpful to gain outputs with higher spatial resolution. Another major obstacle that may prevent the wide application of TISR methods such as DPA-TISR is the requirement on the training dataset. Acquiring a large amount of high-quality low-to-high resolution time-lapse image pairs is laborious and difficult especially when the biological specimen is highly dynamic or of weak fluorescent efficiency. Therefore, further developments of self-supervised or unsupervised learning scheme^3^ for TISR models could greatly reduce their cost for usage. We expected the BioTISR dataset along with the evaluation and development of TISR models would spark more explorations on TISR techniques in the flourishing SR imaging community.

## Methods

### Network configurations for TISR model evaluation

For fair comparison, we designed a template TISR model with the propagation and alignment modules replaceable, as is depicted in Extended Data Fig. 2. The template TISR model was modified from the foundational BasicVSR++ neural network^21^. It is comprised of three primary modules: a feature extraction module, a propagation and alignment module, and a reconstruction module. This architecture is meticulously crafted to be compatible with distinct components of the computational process. For a given image sequence *X* = {*x*_*i*-*k*_, *x*_*i*-*k*+1_, …, *x*_*i*_, …, *x*_*i*+*k*-1_, *x*_*i*+*k*_}, three identical residual blocks are employed to extract features from each frame. Each residual block is comprised of two convolutional layers, a LeakyReLU activation function, and a skip connection. Prior to passing through the residual blocks (RB), a convolutional layer is applied for channel augmentation. The operation of feature extraction module can be formulated as

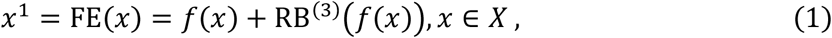

where *f*(·) is a convolutional layer, and

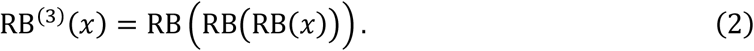

Subsequently, the output feature maps are fed into the propagation and alignment module through two distinct schemes, referred to as sliding window-based propagation (SWP) and recurrent network-based propagation (RNP).

In the SWP scheme, feature maps from the central and adjacent frames within a local window are severally fed into an alignment block (AB) in order to refine all neighbor features towards the central frame. The AB consists of an alignment module, a convolutional layer and three residual blocks. In our validation, three distinct alignment modules based on optical flow, deformable convolution, and non-local attention, respectively, are employed (Extended Data Fig. 2c-e). Afterwards, the resulting feature maps are concatenated, and go through a 1 x 1 convolutional layer to reduce the channels. The progress of SWP can be articulated as follows:

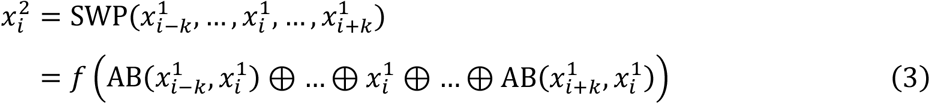

where ⊕ denotes channel-wise concatenation.

In the RNP scheme, the feature maps from adjacent frames generated by the feature extraction module propagate forward and backward in the time dimension following the second-order grid propagation manner^21^. Given the feature map *x*^1^, corresponding features propagated from the first-order neighborhood *h*_*i*+1_, *h*_*i*-1_, and second-order neighborhood *h*_*i*+2_, *h*_*i*-2_, we have

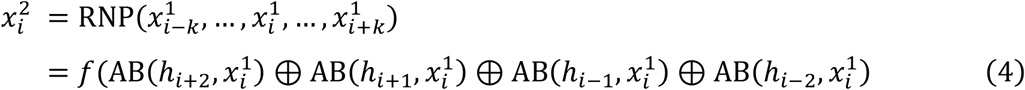

In the optical flow-based alignment block, we compute the optical flow between neighbor frames via the SPyNet^46^. If we denote the optical flow from *x*_*i-k*_ to *x*_*i*_ as *f*_*i-k*→*i*_, the neighbor features *h*_*i-k*_ will be aligned according to *x*_*i*_ by directly warping using the optical flow computed from *x*_*i-k*_ and *x*_*i*_:

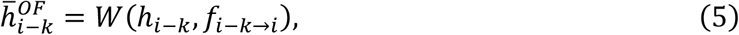

where *W* denotes the spatial warping operation.

In deformable convolution-based alignment block, the optical flow *f*_*i-k→i*_ is used to pre-align the features. The aligned features are then concatenated with the current features 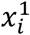 and optical flow *f*_*i-k→i*_ to compute the DC offset *o*_*i-k→i*_ and modulation masks *m*_*i-k→i*_ with two sequential residual blocks:

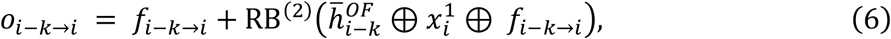

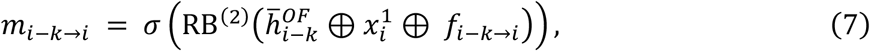

where *σ* denotes the sigmoid activation function. A deformable convolution layer is then applied to the unwrapped feature *h*_*i-k*_:

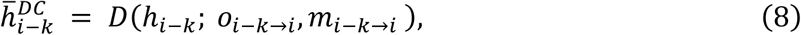

where *D* denotes a deformable convolution.

In non-local attention-based alignment block, three fully connected layers are initially used to generate the embedded query, key and value:

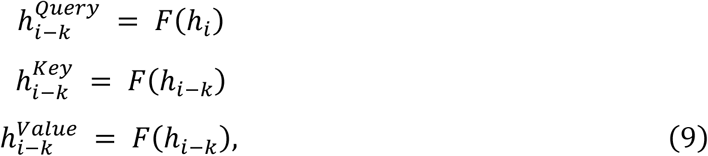

where *F* denotes a fully connected layer. Then the non-local attention mechanism is applied for feature alignment:

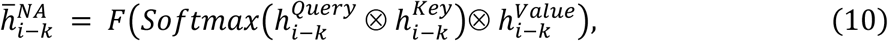

where ⊗ refers to dot product operation. In practice, the embedded features have half of the channel compared to the original feature and we adopt a neighborhood attention mechanism^47^, which can be regarded as a more efficient implementation of nonlocal attention in our experiments, achieving similar SR performance.

Finally, the aligned hierarchical features undergo through the reconstruction module (RM) consisting of three residual blocks, a pixel shuffle layer, and a concluding convolutional layer to generate the SR residuals, which are then added up with up-sampled raw input images, yielding final TISR images. The operation of RM can be formulated as:

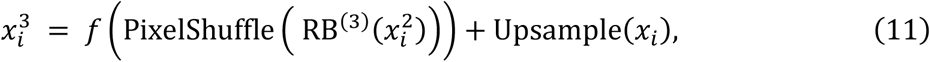

Taking the final output of RM as *ŷ*_*i*_ and the ground truth as *y*_*i*_ for the *i*^*th*^ frame, the overall objective function can be formulated as

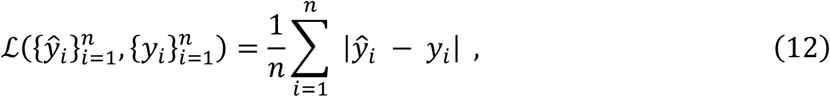

### Network architecture of DPA-TISR

DPA-TISR was constructed based on the optimal TISR baseline model, i.e., adopting recurrent network for propagation and deformable convolution for alignment, following the evaluation conclusions in Fig. 1. In DPA as depicted in Fig. 3, the phase-space alignment mechanism serves as a complementary module to spatial deformable convolution, which synergistically improve the alignment between neighborhood feature maps *h*_*i-k*_ and current feature maps *h*_*i*_. Specifically, the DPA begins with the real-valued fast Fourier transform (denoted as FFT(·), implemented with *torch*.*fft*.*rfft*) to extract the amplitude and phase of both *h*_*i-k*_ and *h*_*i*_:

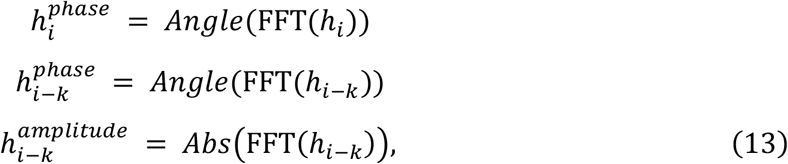

where *Angle*(·) and *Abs*(·) represent the operation to obtain element-wise angle and absolute value of the features. Then the concatenation of phases from current and neighborhood feature maps further goes through a phase-space convolution module, containing one convolutional layer followed by two residual blocks and a skip connection:

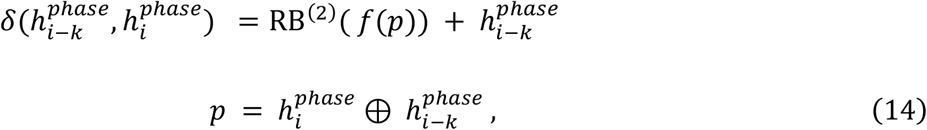

An inverse real-valued fast Fourier transform (denoted as iFFT(·), implemented with *torch*.*fft*.*irfft*) is then utilized to reconstruct the space-domain feature maps from the obtained amplitude and phase components:

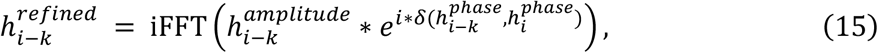

The phase-refined feature maps are then fed into the deformable convolution module as is depicted in Extended Data Fig. 2 for subsequent spatial alignment. The overall architecture of DPA-TISR is illustrated in Extended Data Fig. 5.

### Assessment metrics calculation

We performed image normalization for GT-SIM and TISR images following a commonly used procedure^5, 11^. Specifically, each GT-SIM image stack Y was normalized by dividing by the maximum value, then blurred by a 3×3 size Gaussian kernel with the standard deviation σ = 0.4 (denoted as *ρ*(·)) to mitigate the SIM reconstruction artifacts:

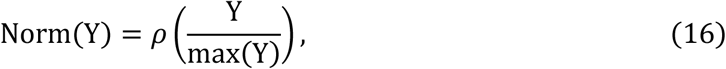

Before computing the assessment metrics, i.e., PSNR and SSIM, a linear transformation was applied to each SR image stack H:

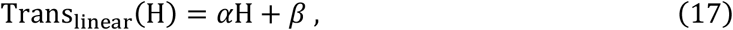

where α and β were chosen by solving the convex optimization problem:

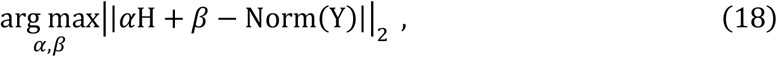

where || · ||_2_ denotes L2-norm. The optimized *α* and *β* result in an MSE-minimized linear transformation of H, effectively scaling and translating every pixel to match the dynamic range of the ground truth.

Three types of metrics were used for quantitatively evaluating the performance in output fidelity, resolution, and temporal consistency, respectively. PSNR and SSIM were utilized to evaluate pixel-level similarity between the inferred SR images and GT-SIM images. Decorrelation analysis^48^ were applied to quantify the image resolution. For temporal consistency assessment, time-lapse Pearson’s correlation matrix was used to visualize the similarity between adjacent SR images 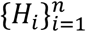. The Pearson correlation between image x and y is calculated by:

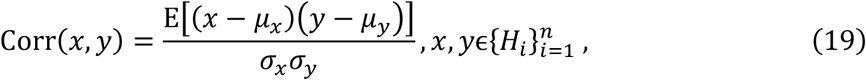

where *µ* and *σ* denote the mean value and standard deviation of corresponding images, respectively; and E represents the arithmetic mean.

### Comparison of DPA-TISR with other models

For the comparative analysis between TISR and SISR models in Figs. 2 and 4, we modified the DPA-TISR into its SISR counterpart by excluding all DPA blocks while keeping other network components, i.e., the feature extraction module and reconstruction module, identical to DPA-TISR. As such, the temporal feature aggregation capability of DPA-TISR was dismissed, thus yielding a SISR manner without too much trainable parameter reduction (∼7M for DPA-TISR versus ∼5.5M for SISR).

In the comparative analysis between DPA-TISR and other TISR models, i.e., VRT and BasicVSR++, we utilized their publicly available implementations on Github^21, 27^. All networks were trained with the same dataset and configurations including the initial learning rate, learning rate decay, batch size, etc., for fair comparison. It’s noteworthy that the patch size of VRT was adjusted to 64, half of that for other models, to ensure similar GPU memory utilization.

### Training details of TISR models

For each type of specimen, we acquired a minimum of 50 groups of WF sequences (512×512 pixels), along with corresponding GT-SIM sequences (1024×1024 pixels for linear SIM and 1536×1536 for nonlinear SIM) in the BioTISR dataset. Typically, we selected ∼35 groups of original data for training and validation, and used the remaining ∼15 groups for testing. Before training, each group of data was augmented into time-lapse image pairs with the size of 128×128×7 for WF input and 256×256×7/384×384×7 for corresponding GT-SIM images by random cropping, horizontal/vertical flipping, and random rotation for further enrichment and avoiding overfitting. In particular, we conducted an evaluation on DPA-TISR models trained with different lengths of input sequence, and found that the input length of 7 is the optimal choice to balance the computation efficiency and SR performance (Supplementary Fig. 8).

The training and inference were performed on a computer workstation equipped with four GeForce RTX 3090 graphic processing cards (NVIDIA) with python 3.6 and PyTorch 1.12.1. In the training phase, the batch size for all experiments was set to 3, and all models were trained using the Adam optimizer with an initial learning rate of 5 x 10^−5^ which was decayed by 0.5 for every 1000 epochs. Training process typically took 18 hours within approximately 3000 epochs. Once the networks were trained, TISR models typically took about 1s to reconstruct a 7-frame SR stack of 1,024×1,024 pixels. The time required for both training and inference decreases linearly with the increase in the number of GPUs utilized. Multi-GPU acceleration has been incorporated into our publicly available Python codes.

### Calculation of uncertainty

The reliability of DPA-TISR predictions can be quantified through two types of uncertainties: model uncertainty and data uncertainty, or referred to as epistemic uncertainty and aleatoric uncertainty, respectively, in Bayesian analysis^16^.

Considering the Non-Bayesian DPA-TISR, represented by *f*_0_, where *θ* is the trainable parameters, the output SR image sequence is denoted as 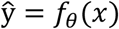. The network parameters are chosen by minimizing the pixel-wise distance between ground truth y and *ŷ*. Taking the L1-loss for example, the objective function can be expressed as follows:

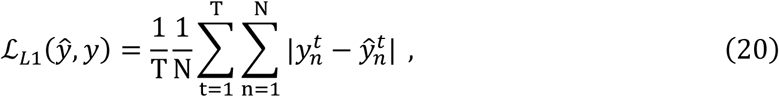

where T denotes the number of timepoints and N denotes the number of pixels in a single image.

Inspired by previous work^11, 17^, we designed the Bayesian DPA-TISR that predicts both the intensity 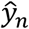 and the scale 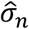 for every pixel. Instead of considering each pixel as a single intensity value, we modeled it as a Laplace distribution empirically:

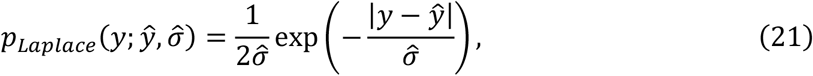

In this way, the scale 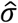 can be regarded as a measurement of the data uncertainty. Then the output SR image 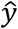 and the scale 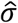 can be simultaneously addressed by minimizing the negative log-likelihood (NLL) function:

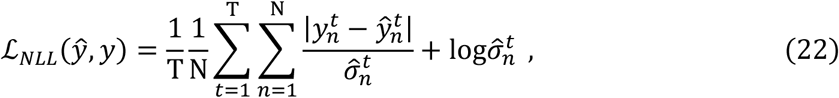

To characterize the model uncertainty, we adopted a Bayesian approximation approach^17^ that employs a distribution over model parameters by incorporating concrete dropout after convolutional layers. Within each inference, the dropout layers in Bayesian DPA-TISR randomly zeroized half of the neurons thus yielding a distinctive network model. Then by aggregating a certain number (denoted as M) of the outcomes of the stochastic forward propagation, the SR results of Bayesian DPA-TISR can be obtained by averaging the predicted intensity of each trial:

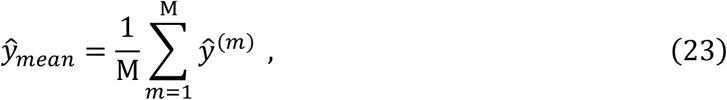

where 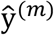 is the predicted intensity map from the *m*^th^ network. The model uncertainty is then quantified by calculated the standard deviation of the predicted results:

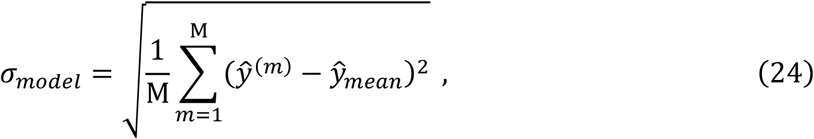

Subsequently, the overall data uncertainty is assessed as follows:

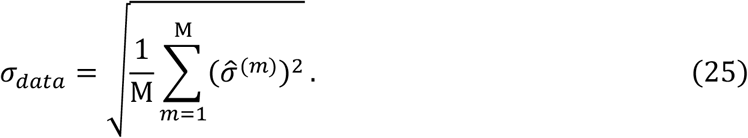

where 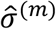 is the predicted scale map from the *m*^th^ network.

### Calculation of confidence map

To enhance the integration of model and data uncertainty information and provide biologists with a more intuitive measurement of uncertainty, we take one step further by employing a pixel-wise mixture probability distribution to generate an integrated confidence map.

During the inference stage, we independently generated M models *θ*^(*m*)^, differing only through dropout layers as depicted above. Considering inferences from M independent models, each pixel *i* follows a mixed Laplace distribution (see Fig. 2a for an example):

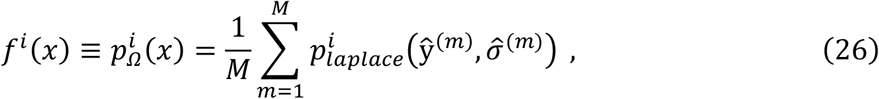

with 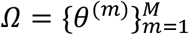, and *x* is the coordinate of the probability distribution function (PDF). Accordingly, we defined the credible interval 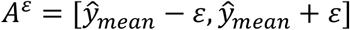 and the interval length *ε*. Then the corresponding confidence *p*^*ε,i*^ for pixel *i* is defined as the probability that the true value *y* falls within *A*^*ε*^:

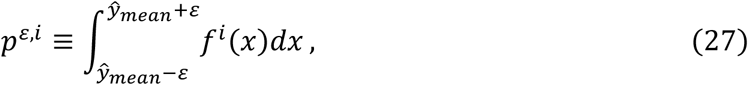

After defining a proper value for ε, which is empirically set as 0.04 by default in our experiments, a confidence map can be generated to indicate the probability that the true value falls within the predicted credible interval.

### Training and inference details of Bayesian DPA-TISR

Based on uncertainty and confidence calculation mentioned above, we designed Bayesian DPA-TISR, which differs from DPA-TISR in three main aspects. First, we integrated dropout layer after each convolutional layer in the feature extraction module and reconstruction module, randomly deactivating neurons during both training and prediction stages. Second, we adjusted the output channel number of the last convolutional layer in DPA-TISR from one to two, representing the predicted mean and scale, respectively. An additional sigmoid function was applied to the predicted scale channel to ensure its non-negativity. Third, the objective function was modified from L1-loss to NLL-loss as formulated in Eq. (22).

During inference stage, we independently executed the trained network with dropout for 6 times, i.e., M=6, averaged the results as the final SR output, and generated confidence map as described previously. Particularly, in long-term confidence-quantifiable TISR imaging experiments, we observed that the predicted scales tended to increase with the corresponding inferred intensities, making it unintuitive to discern which areas of an output image are reliable or unreliable solely from the confidence map. To address this issue, we adopted an intensity-aware confidence generation scheme. Instead of using a constant *ε* of the credible interval 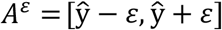, we modified the interval length *ε* as:

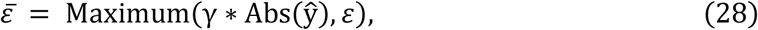

where the scalar scaling factor γ and *ε* were empirically set as 0.2 and 0.04, respectively, in our experiments. Using of 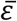 is essentially to define a threshold of 0.2 for foreground segmentation. For pixels with values greater than 0.2, which are more likely to belong to the actual structure, we assigned a proportion (0.2) of their absolute intensity as the interval length. Conversely, for pixels with values less than 0.2, typically representing the background region, we assigned a constant value of 0.04 as the interval length, thereby rationalizing the confidence calculation within background regions of the inferred TISR image.

### Calculation of ECE and reliability diagram

To evaluate the effectiveness of the uncertainty and confidence map, the reliability diagram which compares the empirical accuracy with averaged confidence is computed following a standard procedure^11^ in our experiments. Well-calibrated confidence in the reliability diagram should yield confidence values similar to accuracy, resulting in a diagonal diagram.

Specifically, the empirical accuracy is defined as the proportionality that the ground truth *y* falls into the credible interval 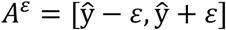. Consequently, the accuracy and confidence are specified as follows:

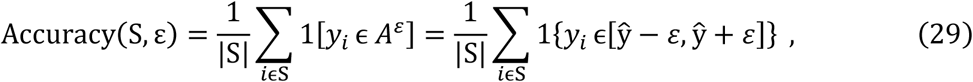

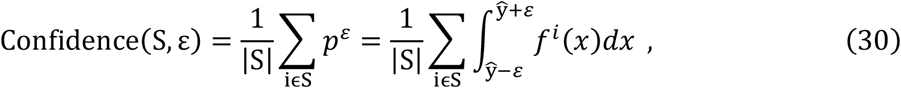

where S denotes a subset of all pixels, 1(·) denotes the indicator function, and ε > 0 determines the length of the credible interval around each 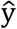.

To construct a reliability diagram with K groups, we divided the value of confidence into K intervals segmented by *τ*_0_, *τ*_1_, …, *τ*_*K*_. For pixels in 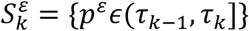, we plotted Confidence 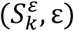 against Accuracy 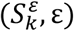 to obtain the final reliability diagram. Furthermore, the expected calibration error (ECE) is determined as the weighted average of the absolute differences between the accuracy and confidence:

### Confidence correction for Bayesian DPA-TISR

Recognizing the disparity between the estimated confidence and actual accuracy, i.e., over confidence for most Bayesian neural networks, we developed an iterative finetuning framework to eliminate the ECE between the estimated confidence and accuracy. During the finetuning stage, the objective function is modified from Eq. (22) as follows:

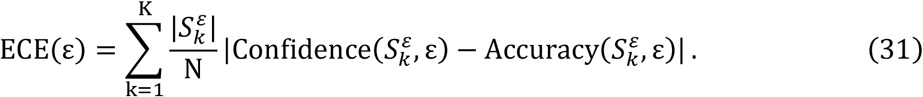

where *R*_*confidence*_ is the confidence correction regularization (CCR), and *α* is a weighting scalar to balance ℒ_*NLL*_ and *R*_*confidence*_, which is empirically set as 0.1 in our experiments. CCR is comprised of two parts:

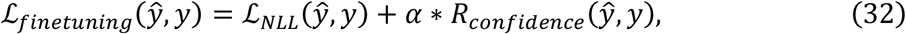

where the second term, *confidence*(*x*), is the average confidence of the outputs and the corresponding weighting scalar *k* aims at adjusting overall average confidence of prediction. Positive values of *k* suppress over-confidence and negative values of *k* restrain under-confidence, which is the key variable in our optimization framework. We adopted a combined strategy of linear searching and parabola fitting to determine the optimal value of *k*.

In the optimization process, a relative ECE (rECE) of the original trained network, denoted as *E*_0_, is first calculated by

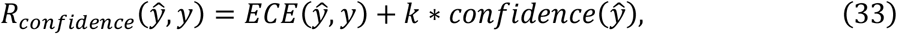

of which the sign is used to determine the initial directions for subsequent optimization:

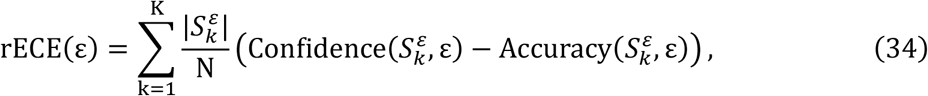

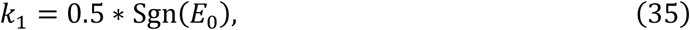

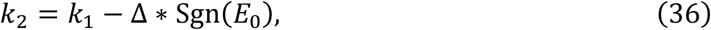

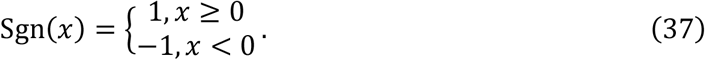

where Δ is the step size set as 0.1 by default. Next, the first two trials of finetuning are independently performed with *k*_1_and *k*_2_, respectively, using the objective function described in Eq. (32), after which, we obtained two new ECE values of the finetuned networks, denoted as *E*_1_ and *E*_2_. If *E*_1_ < *E*_2_, we exchange *k*_1_ and *k*_2_ as well as the corresponding *E*_1_ and *E*_2_, and reverse the searching direction, i.e., reset Δ as -0.1, to ensure *k*_2_ is the ECE-descent direction compared to *k*_1_. Afterwards, the linear searching process continues along this descent direction

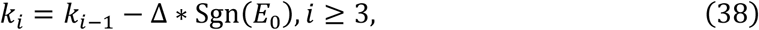

until finding a *k*_*i*_ that satisfies *E*_*i*_ > *E*_*i*-1_, where *E*_*i*_ denotes the ECE value of the finetuned model with *k*_*i*_. We then utilized the quadratic polynomial fitting to find the optimal *k*_*_ value according to three latest weighting scalars *k*_*i*_, *k*_*i*-1_, *k*_*i*-2_ and corresponding ECE values *E*_*i*_, *E*_*i*-1_, *E*_*i*-2_ :

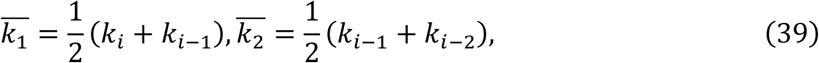

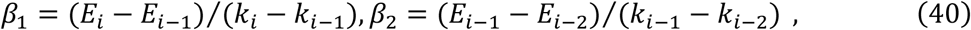

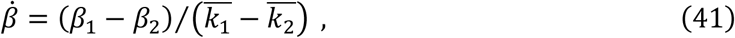

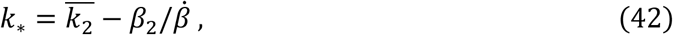

Finally, one last finetuning is carried out using the optimal *k*_*_ in the objective function to obtain a confidence-calibrated Bayesian DPA-TISR model. During the fine-tuning stage, the models were trained using the Adam optimizer with an initial learning rate of 5 × 10^−5^, which is decayed following a cosine annealing strategy over a course of 30 epochs. The overall finetuning process typically takes less than 1 hour. The workflow diagram of the confidence correction is shown in Supplementary Fig. 9.

### Mito-PO contact quantification

The contact level between Mito and POs was evaluated by confidence-weighted Mander’s overlap coefficient (MOC). Considering that Mito-PO contact sites were principally situated at the periphery of POs, we first identified the boundary of each PO using the following procedure: (i) Estimate the background of the region of interest (ROI) of the PO by applying a Gaussian filter with a standard deviation σ = 10 pixels; (ii) Smooth the TISR image using another Gaussian filter with σ = 1 pixel and then subtract the estimated background; (iii) Generate a binary mask by setting a threshold to the background-subtracted image; (iv) Extract the boundary of each PO using the Sobel operator; (v) The PO boundary image is convolved with the equivalent PSF of SIM, thereby delineating potential contact regions of POs. Next, following the aforementioned steps (i)-(iii), a binary Mito mask, denoted as M_Mito_, was calculated.

We reasoned that the regions with higher confidence should have higher weights in MOC calculation. Therefore, we applied the confidence map estimated by the Bayesian DPA-TISR model as adaptive wights to rationalize the quantification of Mito-PO contacts as follow:

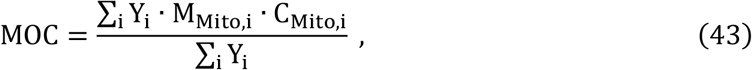

where Y_i_, M_Mito,i_, and C_Mito,i_ denote the value of the *i*^th^ pixel in the ROIs of the PO boundary image Y, Mito mask M_Mito_ and the confidence map C_Mito_.

### Cell culture, transfection, and staining

COS-7 and HeLa cells as well as their stable cell lines were cultured in DMEM (Gibco, cat. no. 11965092), supplemented with 10% fetal bovine serum (Gibco, cat. no. 10099141C) and 1x penicillin-streptomycin (Thermo Fisher, 15140122) under 37 °C in Thermo Scientific™ Heracell™ 150i CO_2_ incubator. SUM159 cells were cultured in DMEM/F12K medium supplementary with 5% Fetal Bovine Serum (FBS) and 1% Penicillin-Streptomycin solution.

For live cell imaging, the 35 mm coverslips were pre-coated with 50μg ml^-1^ of collagen and 1 × 10^5^ cells were seeded onto coverslips. For transient transfection, cells were transfected with plasmids using Lipofectamine 3000 (Invitrogen, cat. no. L3000150) according to the manufacturer’s protocol 12 hours post plating. Cells were imaged for 12 hours after transfection. Where indicated, the cells transfected with Halo Tag plasmids were labelled with 10 nM JF549 ligand for 15 min according to the published protocol^49^. The cells were rinsed with fresh medium to remove unbound ligand and imaged immediately afterward. The plasmids used in transient transfection include Lifeact-mEmerald, Lifeact-SkylanNS, Clathrin-mEmerald, Clathrin-mCherry, 3 xmEmerald-Ensconsin, Lamp1-Halo, 2 xmEmerald-Tomm20, TFAM-mEmerald, PKMO-Halo, and PMP-Halo.

### Statistics and reproducibility

Figs 1h-1j, 2e, 2f, and Extended Data Fig. 3c, 3d, 6f, 6h were plotted in Tukey box-and-whisker format. The box extends from the 25th and 75th percentiles and the line in the middle of the box indicates the median. The upper whisker represents the larger value between the largest data point and the 75th percentiles plus 1.5× the interquartile range (IQR), and the lower whisker represents the smaller value between the smallest data point and the 25th percentiles minus 1.5× the IQR. Data points larger than the upper whisker or smaller than the lower whisker is identified as outliers, which are displayed as black spots.

## Data availability

The BioTISR dataset will be made publicly accessible at the Zenodo repository after the paper is accepted by a peer-reviewed journal. All data that support the findings of this study in Figs. 1-5, Extended Data Figs. 1-10, Supplementary Figs. 1-9, and Supplementary Videos 1-5 are available upon requests.

## Code availability

The python codes of Bayesian DPA-TISR, several representative pre-trained models, as well as some example data for testing are already publicly accessible on Github (https://github.com/liushuran2/Bayesian_DPA_TISR).

## Acknowledgements

The authors thank T. Kirchhausen for the donor plasmids used for genome editing and help in generating the genome-edited cell lines, and thank Dr. Kangmin He for genome-edited SUM159 cell lines. This work was supported by grants from the National Natural Science Foundation of China (32125024, 32271513, 62322110, 62071271, and 62088102); the National Key R&D Program of China (2023YFB3209700); the Ministry of Science and Technology (2021YFA1300303 and 2020AA0105500); the Chinese Academy of Sciences (ZDBS-LY-SM004 and XDA16021401); the Collaborative Research Fund of the Chinese Institute for Brain Research, Beijing (2021-NKX-XM-03); China Postdoctoral Science Foundation (2022M721842, 2023T160365); the New Cornerstone Science Foundation; the Shuimu Tsinghua Scholar Program (2022SM035); Beijing Natural Science Foundation (JQ21012).

## Author contributions

Q.D., D.L., and H.Q. supervised the research. Q.D., D.L., and C.Q. conceived and initiated this project. C.Q., S.L., and Y.W. designed the detailed implementations. S.L. and C.Q. developed the python code, performed simulations, and processed relevant imaging data. T.J., X.G., and W.X. prepared samples and performed imaging experiments. S.L., C.Q., and Q.M. analyzed the data with conceptual advice from Q.D., D.L., and H.Q. C.Q., S.L., W.X. and J.Z. composed the figures and videos under the supervision of Q.D. and D.L. C.Q., D.L, S.L., and Y.W. wrote the manuscript, with input from all authors. All authors discussed the results and commented on the manuscript.

## Supplemental information

Supplemental Information includes 5 notes, 9 figures, and 5 videos.

## Extended Data Figures

**Extended Data Fig. 1.**
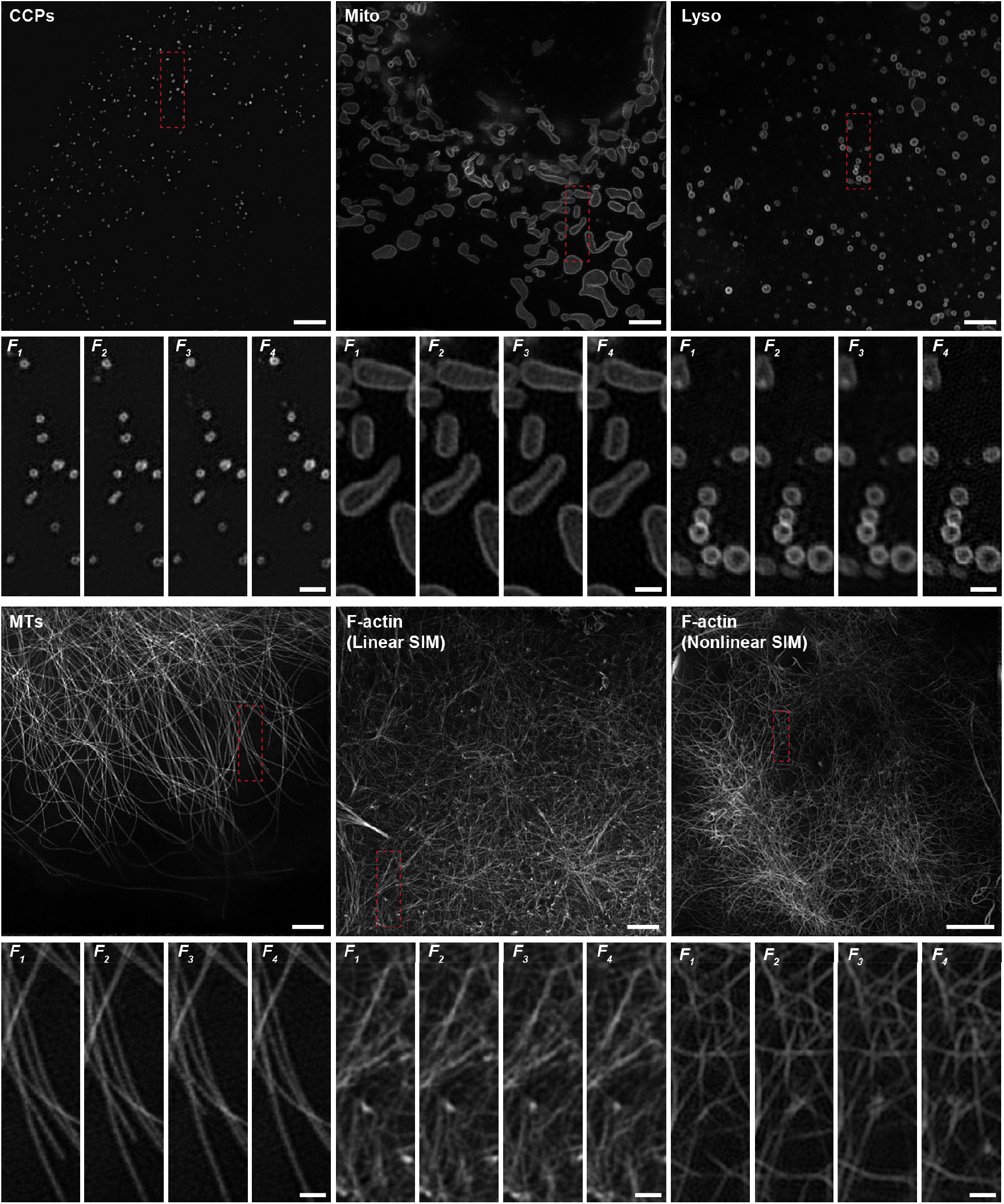
Gallery of BioTISR dataset. Representative time-lapse GT-SIM images of CCPs, Mito, Lyso, MTs, F-actin (linear SIM), and F-actin (nonlinear SIM). Scale bar: 3μm, and 1μm (zoom-in regions). Gamma value: 0.8 for F-actin images of both linear and nonlinear SIM.

**Extended Data Fig. 2.**
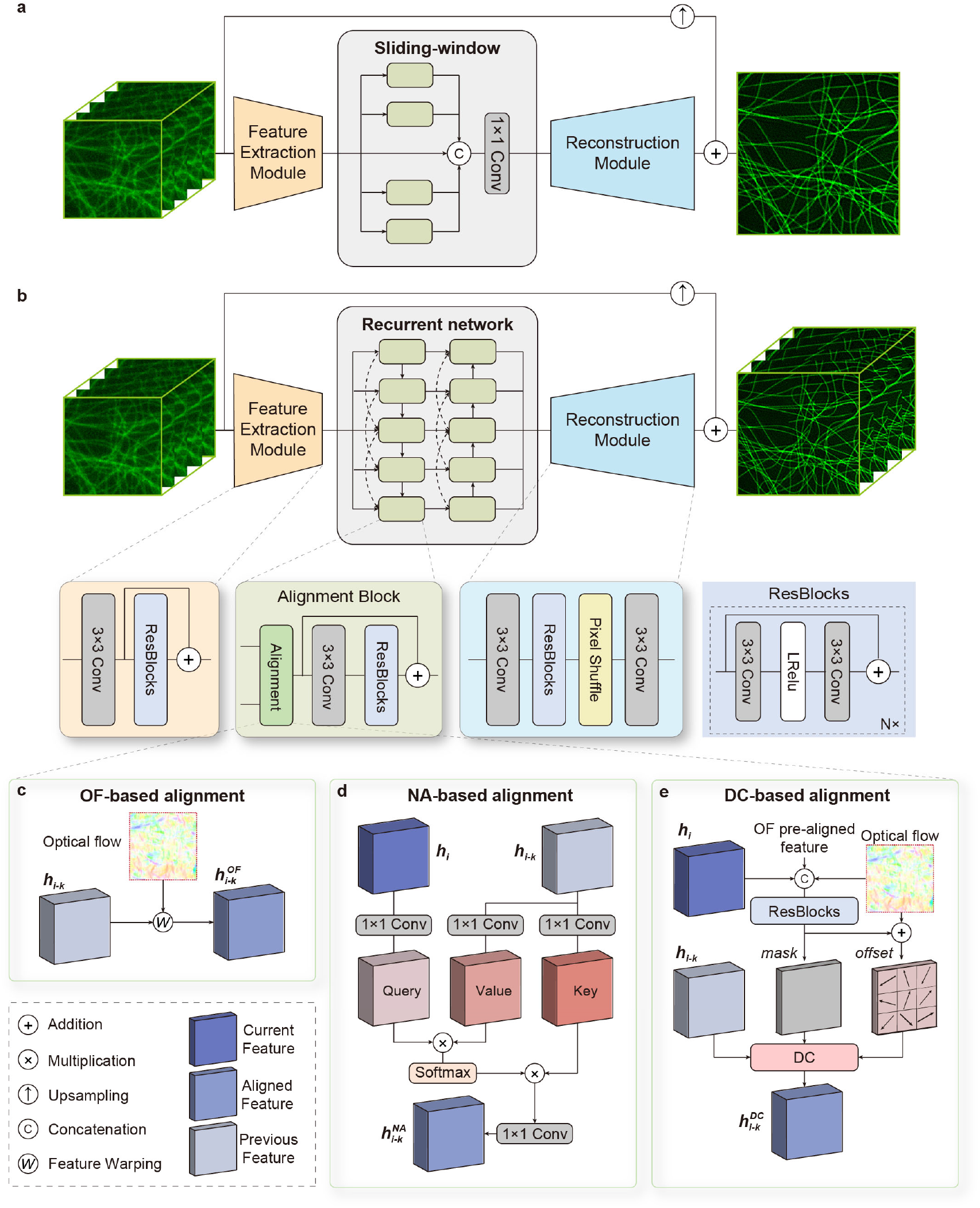
Template TISR neural network architecture used in evaluation. **a**,**b**, Template TISR neural network architectures equipped with sliding window-based propagation (a) and recurrent network-based propagation (b). **c**-**e**, Three representative neighbor feature alignment mechanisms based on optical flow (OF, c), nonlocal attention (NA, d), and deformable convolution (DC, e).

**Extended Data Fig. 3.**
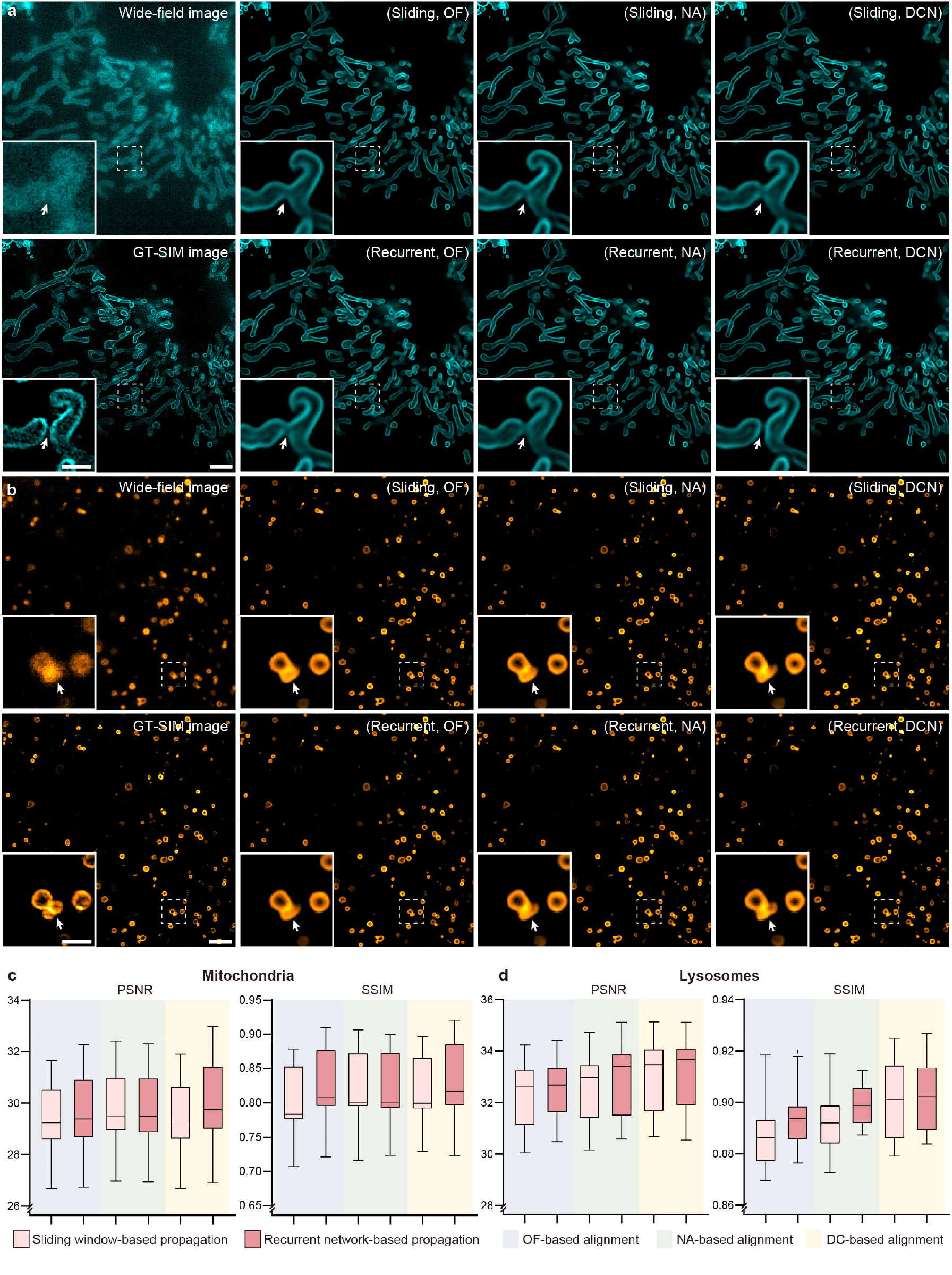
Comparison of representative propagation and alignment mechanisms in TISR models with nonlinear SIM dataset of F-actin. **a**,**b**, Representative TISR images of mitochondria (a) and lysosomes (b) inferred by six models combined by two propagation methods, sliding window-based propagation (simplified as sliding) and recurrent network-based propagation (simplified as recurrent), and three alignment mechanisms based on optical flow (OF), nonlocal attention (NA) and deformable convolution (DC). WF and GT-SIM images are shown in the first column for reference. **c**,**d**, Statistical comparison of the six models in terms of PSNR, SSIM on images of mitochondria (c, n=50) and lysosomes (d, n=50). Scale bar, 3 μm (a, b), 0.5 μm (zoom-in regions in a and b).

**Extended Data Fig. 4.**
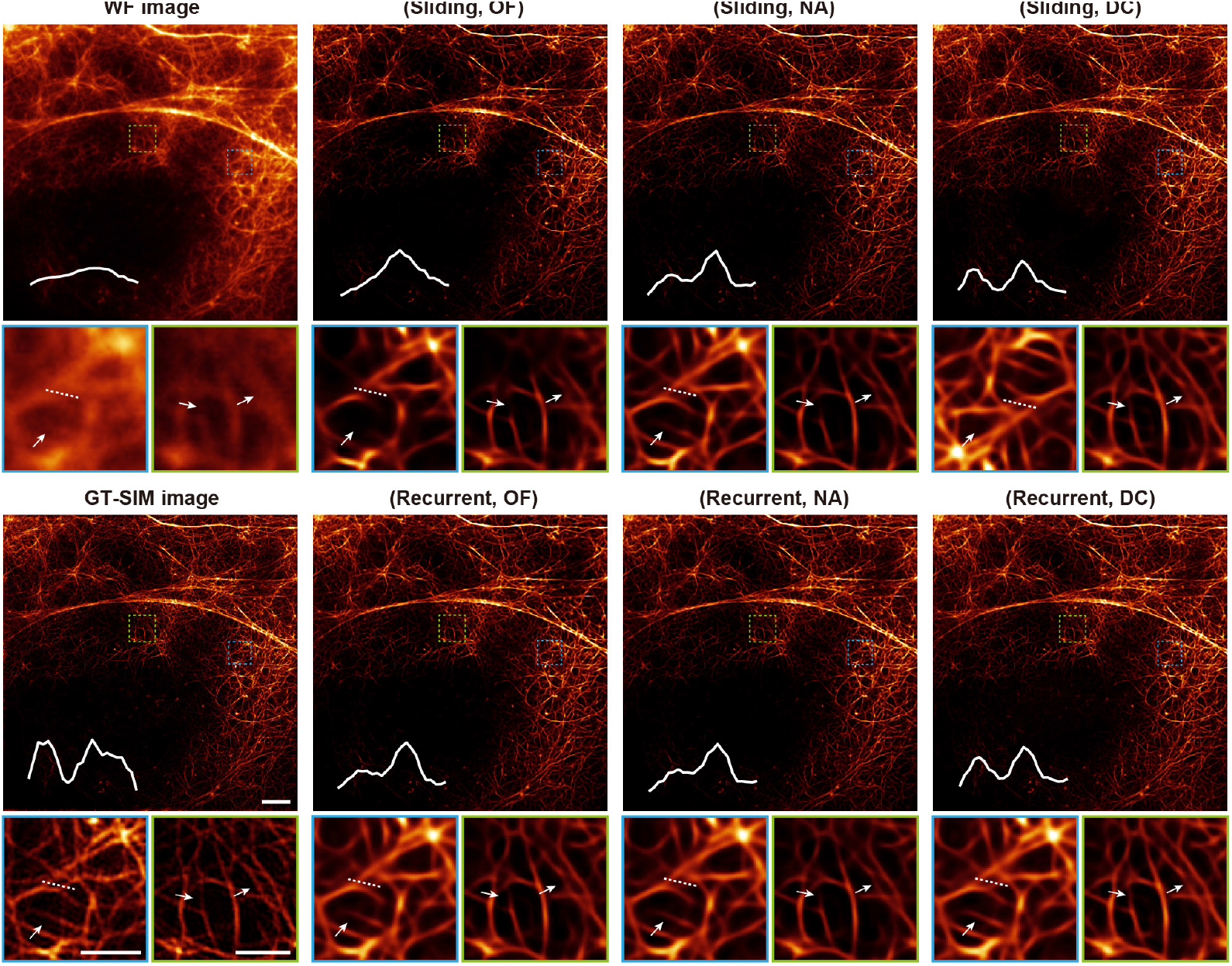
Comparison of representative propagation and alignment mechanisms in TISR models with nonlinear SIM dataset of F-actin. Representative TISR images inferred by six models combined by two propagation methods, sliding window-based propagation and recurrent network-based propagation, and three alignment mechanisms based on optical flow (OF), nonlocal attention (NA) and deformable convolution (DC). All models are configured to up-scaling the input image by 3-fold and trained with the nonlinear SIM F-actin dataset in BioTISR. WF and nonlinear GT-SIM images are shown in the first column for reference. The intensity profiles along the white dotted lines in magnified images are shown on the bottom left corner of each image. Scale bar, 3 μm, 1 μm (zoom-in regions).

**Extended Data Fig. 5.**
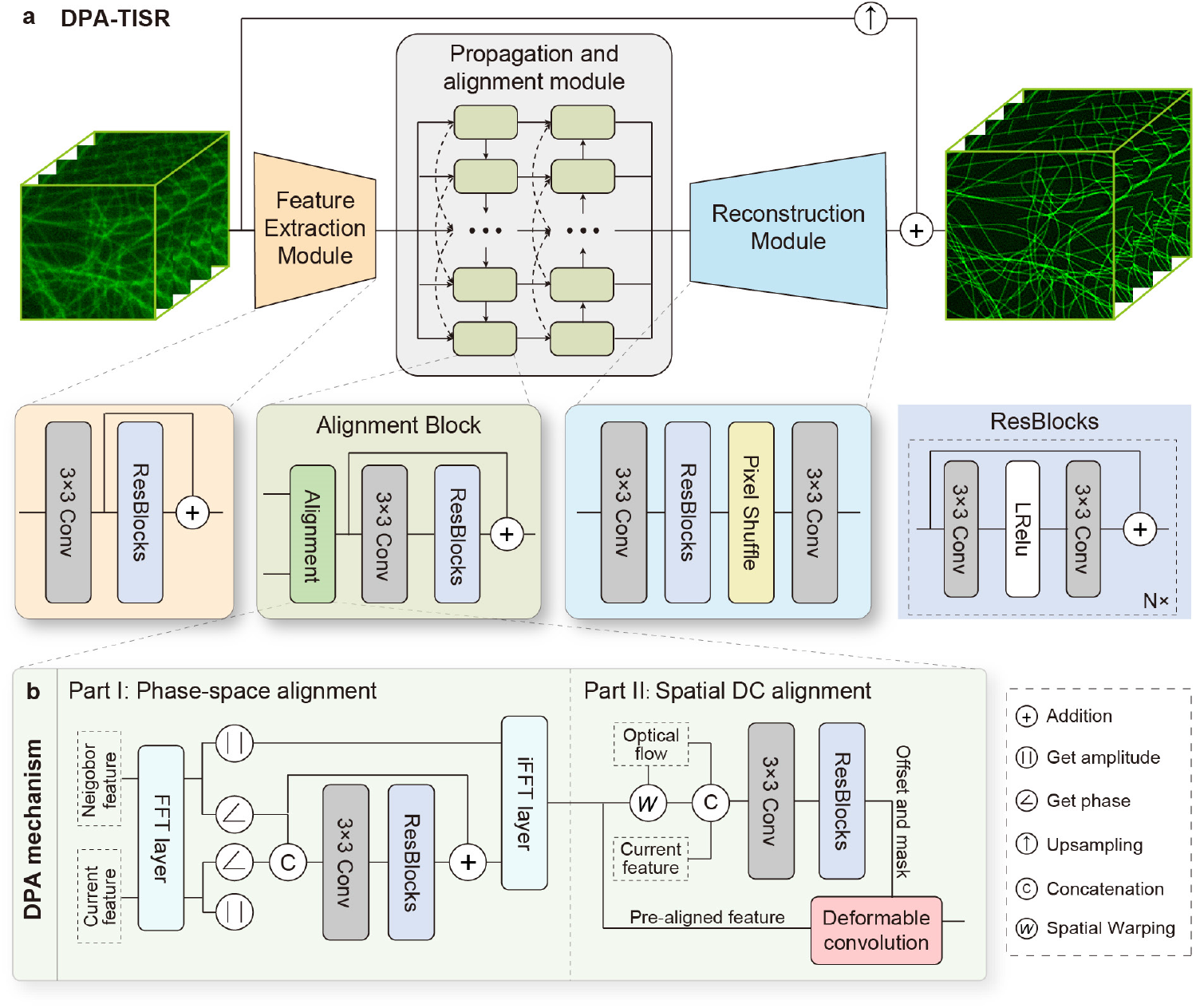
Network architecture of DPA-TISR. **a**, Overall network architecture of DPA-TISR model, which consists of the feature extraction module, propagation and alignment module, and reconstruction module. **b**, Deformable phase-space alignment (DPA) mechanism which is sequentially comprised of two parts: the phase-space alignment (left panel) and spatial deformable convolution (DC) alignment (right panel).

**Extended Data Fig. 6.**
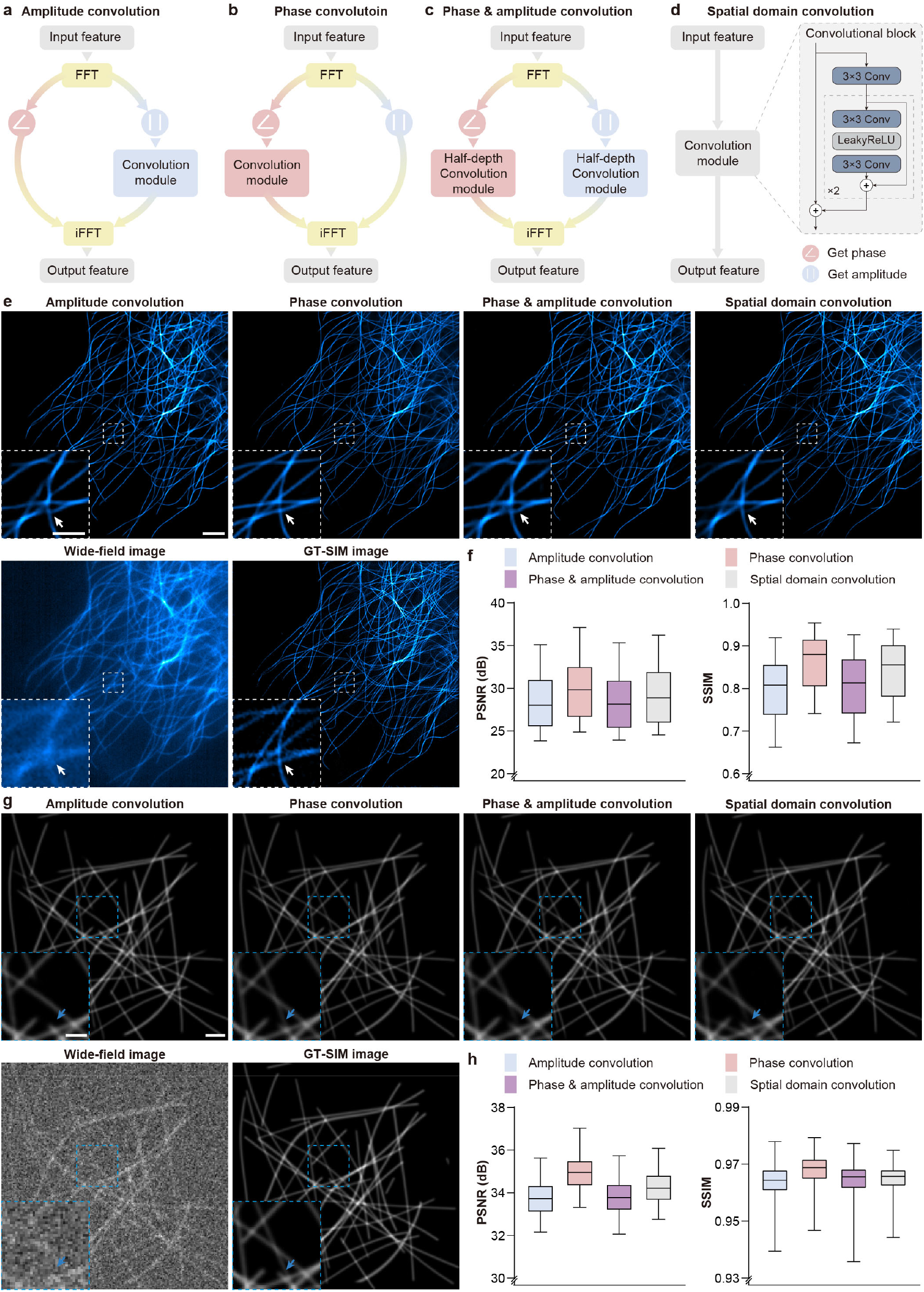
Comparison of the phase-space alignment mechanism and its variants. **a-d**, Four variants of phase-space alignment in which convolution is imposed on the amplitude (a), phase (b), both amplitude and phase each with half of the convolutional depth in the frequential space (c) and pure spatial features (d). TISR models compared in here are identical except for the pre-alignment methods in the alignment block (Supplementary Note 3). **e**, Representative TISR images of microtubules inferred by models equipped with four different pre-alignment mechanisms depicted in a-d. The input wide-field image and GT-SIM image are provided for reference. **f**, Statistical comparison in terms of PSNR and SSIM for microtubule SR images inferred by TISR models with four alignment variants (n=50). **g**, Representative TISR images of simulated tubular structures inferred by models equipped with four different pre-alignment mechanisms depicted in a-d. The input wide-field image and GT-SIM images are provided for reference. **h**, Statistical comparison in terms of PSNR and SSIM for simulated images inferred by TISR models with four alignment variants (n=200). These results demonstrated that feature alignment in frequential space generally outperforms alignment in purely spatial domain. Among three frequential alignment mechanisms, phase convolution-based alignment surpasses other configurations with similar convolutional depth and computation complexity, indicating the superiority of the proposed phase-space alignment mechanism. Scale bar, 3μm (e, g), 1μm (zoom-in regions of e and g).

**Extended Data Fig. 7.**
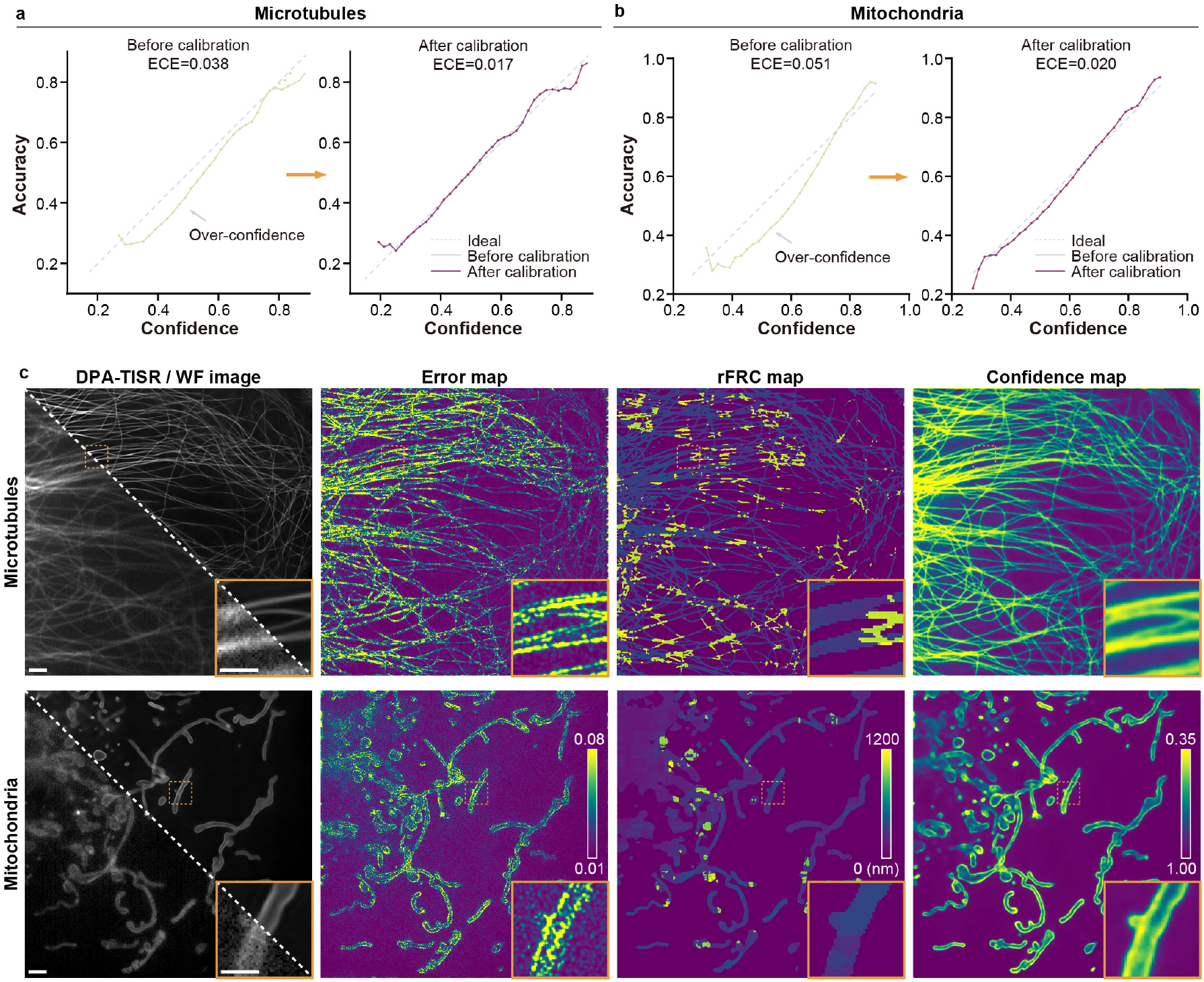
Confidence calibration of Bayesian DPA-TISR models trained with microtubule and mitochondrion datasets. **a**,**b**, Reliability diagrams generated by Bayesian DPA-TISR models before (left panel) and after (right panel) confidence calibration for microtubule (a) and mitochondrion images (b). **c**, Representative wide-field (WF) images (bottom left corner of the first column), TISR images (top right corner of the first column), absolute error maps (second column), rFRC maps generated with rolling Fourier ring correlation^36^ (third column), and confidence map estimated by Bayesian DPA-TISR models (four column) of microtubules and mitochondria. Scale bar: 1μm (c), and 0.5μm (zoom-in regions in c).

**Extended Data Fig. 8.**
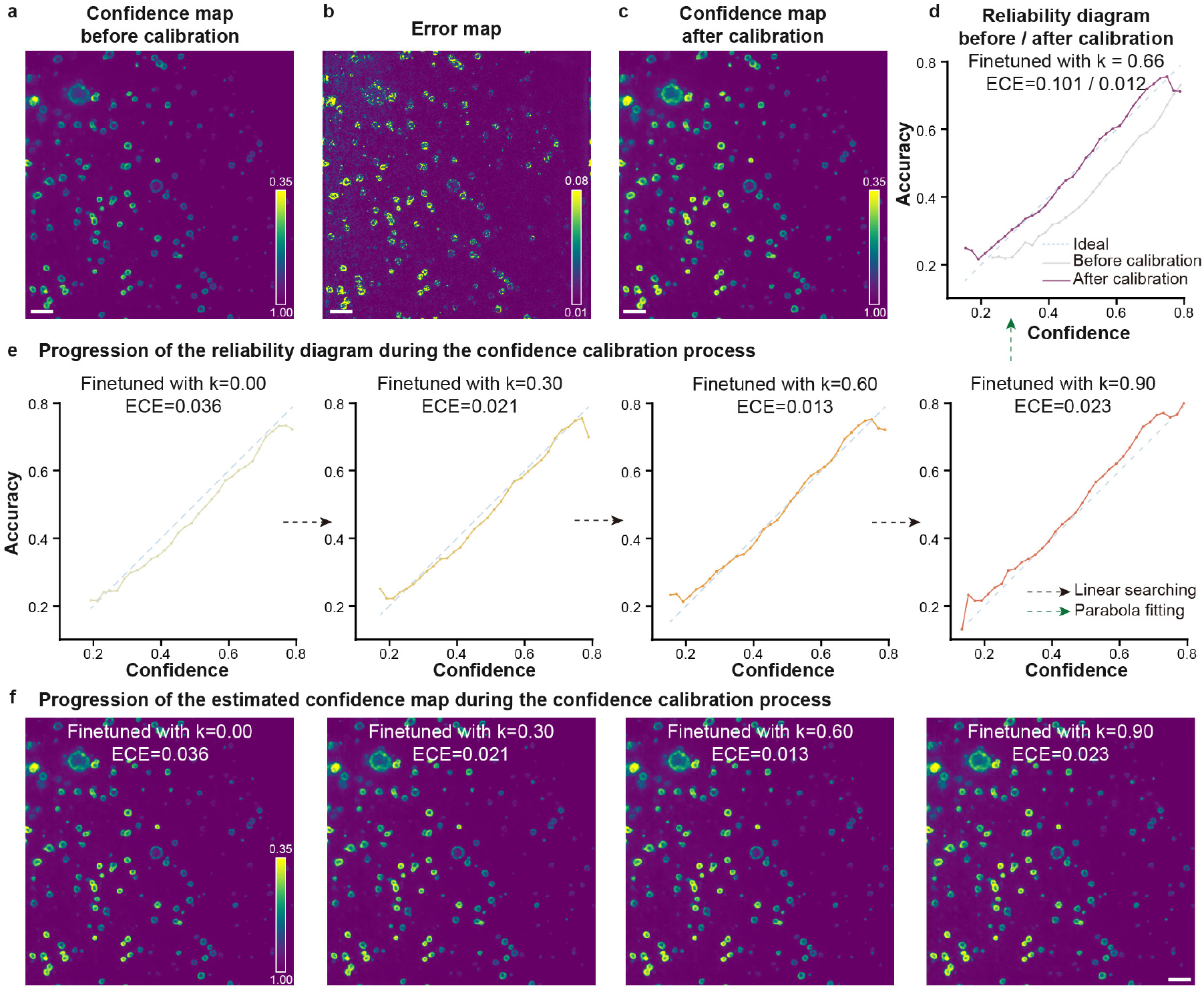
Confidence calibration for the Bayesian DPA-TISR model trained with lysosome images. **a**, Confidence map generated by the Bayesian DPA-TISR model before confidence calibration. **b**, The absolute error map calculated with the inferred TISR image and GT-SIM image shown for reference. **c**, Confidence map generated by the Bayesian DPA-TISR model after confidence calibration. **d**, Reliability diagrams presented by accuracy versus average confidence before and after confidence correction with corresponding expected calibration error (ECE) and *k* value labeled. **e**,**f**, Representative detailed searching steps of the iterative finetuning process. The reliability diagrams (e) and corresponding confidence maps (f) are shown. Scale bar, 1 μm (a-c and f).

**Extended Data Fig. 9.**
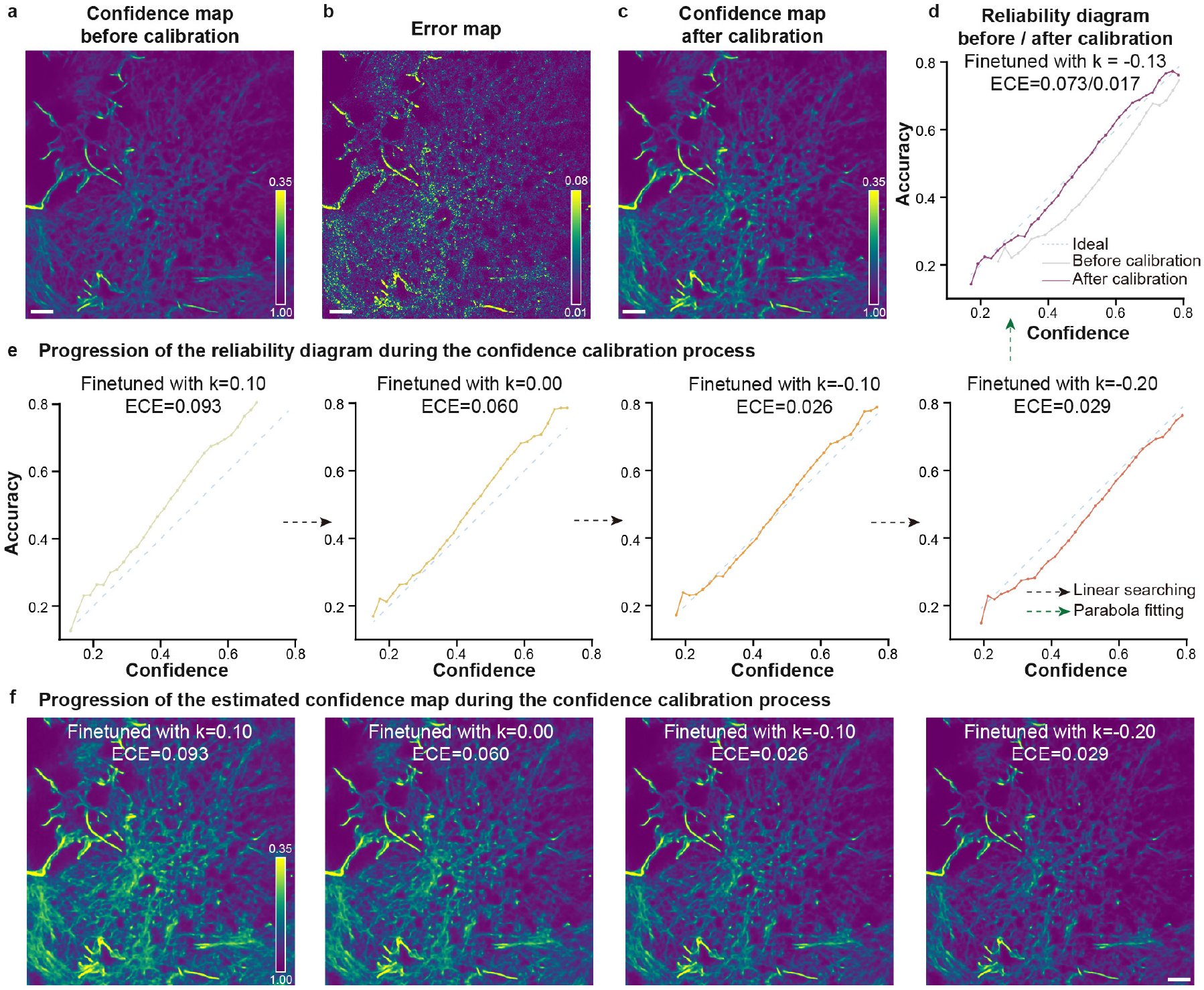
Confidence calibration for the Bayesian DPA-TISR model trained with F-actin images. **a**, Confidence map generated by the Bayesian DPA-TISR model before confidence calibration. **b**, The absolute error map calculated with the inferred TISR image and GT-SIM image shown for reference. **c**, Confidence map generated by the Bayesian DPA-TISR model after confidence calibration. **d**, Reliability diagrams presented by accuracy versus average confidence before and after confidence correction with corresponding expected calibration error (ECE) and *k* value labeled. **e**,**f**, Representative detailed searching steps of the iterative finetuning process. The reliability diagrams (e) and corresponding confidence maps (f) are shown. Scale bar, 1 μm (a-c and f).

**Extended Data Fig. 10.**
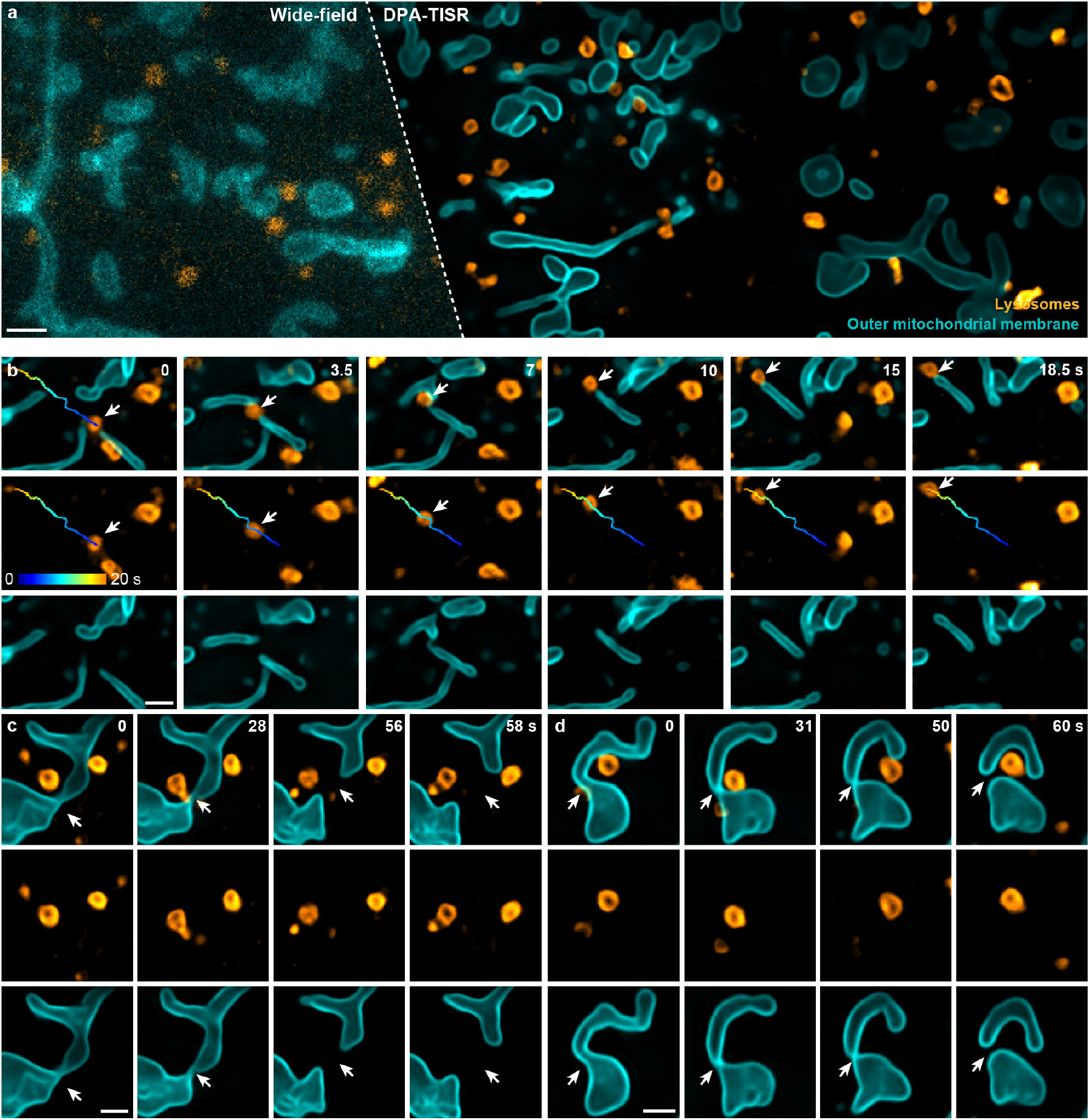
Long-term SR live imaging of interactions between mitochondria and lysosomes enabled by DPA-TISR. **a**, Representative two-color DPA-TISR frame of outer mitochondrial membrane and lysosomes from a long-term video of ∼10,000 timepoints. Wide-field image before processing is shown in the left panel for comparison. **b**, Time-lapse TISR images showing that a mitochondrion underwent the directional movement by hitchhiking on a moving lysosome. The temporally color-coded trajectory of the lysosome is plotted. **c**,**d**, Time-lapse images visualizing two typical cases of mitochondrial fission under the mediation of lysosomes at the contact sites. Scale bar: 3μm (a), and 1μm (b-d).

