## Supplementary Materials for "Time-lapse Image Super-resolution Neural Network with Reliable Confidence Quantification for Optical Microscopy"

**This PDF file includes:**

Supplementary Notes 1-5

Supplementary Figures 1-9

Captions for Supplementary Videos 1-5

Supplementary References

### Supplementary Notes

#### 1. General consideration of TISR models

The resolution of an optical microscope is physically constrained by the diffraction limit, leading to a finite-size point spread function (PSF) in the spatial domain. This inherent limitation implies that a conventional optical microscope cannot distinguish structures finer than ~200 nm, even with the higher magnification or spatial sampling rate beyond the Nyquist criterion.

Super-resolution (SR) microscopy has revolutionized imaging capabilities by surpassing the long-standing limitations of diffraction-limited resolution. This breakthrough is achieved through precise modulation of fluorescence excitation or emission in spatially coordinated manners, such as stimulated emission depletion (STED) microscopy[1, 2], structured illumination microscopy (SIM)[3], or in a stochastic excitation manner, such as stochastic optical reconstruction microscopy (STORM)[4] and photoactivated localization microscopy (PALM)[5], followed by computational reconstruction to obtain the final SR images. However, it is noteworthy that the increase in spatial resolution obtained through any hardware SR approach necessitates longer acquisition time and/or higher illumination intensity, compromising other equally important performance, e.g., speed and duration, in live-cell imaging. While SIM has been gaining popularity for SR live-cell imaging with its low phototoxicity and high temporal resolution, its post-reconstruction process requires relatively high signal-to-noise ratios (SNR) for each raw image to produce high-quality SR-SIM images, impairing fast, low-light, and quantitative imaging. Although multiple improvements in all aspects of SR techniques have been exploring in recent years, the fundamental problems of conventional hardware-based super-resolution imaging methods, including the long acquisition time and severe phototoxicity, still hinder their wider application.

On the other hand, the rapid development of artificial intelligence has assisted conventional SR techniques in surpassing many traditional hardware limitations. For example, various deep learning networks have displayed excellent performance in the single-image super-resolution (SISR) task, which usually transforms a single low-resolution (LR) image to a high-resolution (HR) counterpart. Recently, state-of-the-art neural network models or their variants in computer vision have been adapted to enhance the resolution of microscopic images, including cross-modality transformation[6, 7], self-learning-based axial resolution enhancement[8-10], and SR reconstruction from raw images[11-13]. The integration and innovation of artificial intelligence algorithms with conventional microscopy technologies have emerged as a fast-growing interdisciplinary research field, significantly affecting the development of SR microscopy.

Nevertheless, SISR-based computational approaches also pose substantial challenges and limits. First, the inherent quantum nature of photons introduces unavoidable stochasticity in optical measurements. In fluorescent imaging and neural network reconstruction, detection noise exacerbates the uncertainty of the results and hinders the visualization of underlying structures. As is shown in Supplementary Fig. 6, the quality of the network output deteriorated as the SNR of the input wide-field images decreased. Second, previous SISR models neglect the temporal consistency of biological imaging, resulting in large difference between two consecutive frames, thereby disrupting the observation of crucial bioprocesses. This lack of temporal stability further deteriorates long-term imaging quality.

Taken together, we believe that incorporating temporal information is crucial to improving spatial resolution, reconstruction fidelity, and temporal consistency in image super-resolution tasks. When processing time-lapse images, a more effective approach compared to existing SISR methodology is to input multiple consecutive frames simultaneously into the network, which allows information from different frames to propagate through each other, leading to optimal results.

#### 2. Generation of simulated time-lapse images

##### I. Simulation of ground truth images of the tubular structure

We generate the ground truth (GT) images $I_{GT}$ of tubular structures following our previously described procedures[11], which briefly consists of three steps:

(i) Utilize MATLAB's *randi()* function to randomly generate positional parameters for tubules and then employ the *insertShape()* function to sketch them onto a blank image of 384×384 pixels;

(ii) Assign a random elastic deformation field comprised of 4×4 elements and resize it to the same size as the generated image of tubular structure with bicubic interpolation;

(iii) Utilize the MATLAB function *interp2()* to implement the resized elastic deformation field onto the sample image, thereby inducing curvature in the tubular structures.

To mimic the active dynamics of microtubules in live cells and generate corresponding time-lapse image sequence, we simulate a displacement for each simulated tubules between adjacent two frames. Specifically, an initial sample image is generated following the aforementioned procedure. Subsequently, a displacement $D_{i,t}$ is assigned along a randomly specified orientation for each simulated tubule $i$ at each time point $t$ through the MATLAB function *imwarp()*. The displacement $D_{i,t}$ is defined as

$$D_{i,t} = \frac{1 + 2u_{i,t}}{3} v, \tag{1}$$

where $u_{i,t}$ is randomly sampled from the uniform distribution $\mathrm{U}[0,1]$, and $v$ is the pre-defined velocity parameter representing the moving speed of current image sequence, which is set to $\sim 1 \mu m/s$, consistent with the growth velocity of microtubules in live COS-7 cells. The pixel size of the generated sample images is 30.3 nm, which is set to be half the pixel size of our Multi-SIM system.

##### II. Simulation of diffraction-limited wide-field images and GT-SIM images

To simulate the diffraction-limited wide-field (WF) image sequence of tubular structures, each of the ground truth images is convolved with a wide-field point spread function (PSF) based on our imaging configuration of $\mathrm{NA}_{detection} = 1.35$, $\lambda_{emission} = 515\mathrm{nm}$. The PSF is normalized so that the summation of all pixel values is equal to 1. The resulted images are then down-sampled by 2-fold via bicubic interpolation to generate diffraction-limited wide-field images without noise, denoted as $I_{clearWF}$. Finally, the wide-field images contaminated by Poisson noise is generated via:

$$I_{noisyWF} = N_{Poisson}(\alpha I_{clearWF}) + G(0, {\sigma_r}^2) + b_c \tag{2}$$

where, $\alpha$ is the coefficient used to tune the signal level; $b_c$ is the camera background set as 100; $N_{Poisson}$ denotes Poisson corruption, $G(0, {\sigma_r}^2)$ denotes a white noise image with zero mean and variance of ${\sigma_r}^2$ measured from background frames of our sCMOS camera.

On the other hand, the GT-SIM image sequence, $I_{GT-SIM}$, is generated by simply convolving $I_{GT}$ with SIM PSF, which is 2 fold narrower than the wide-field PSF:

$$I_{GT-SIM} = I_{GT} \otimes PSF_{GT-SIM} \tag{3}$$

where $\otimes$ is the convolution operation.

#### 3. The design of DPA-TISR

Compared to SISR, which concentrates on exploiting the intrinsic properties of a single image to recover as much as the high-frequency information, TISR presents an additional challenge that it involves aggregating information from multiple highly-related but misaligned frames in time-lapse inputs. To this end, we firstly examined whether the state-of-the-art (SOTA) deep neural networks (DNNs) designed for natural video super-resolution (VSR), i.e., super-sampling, tasks could serve as viable solutions for TISR models in fluorescence imaging.

Most existing deep neural networks for VSR consist of four interrelated components: propagation, alignment, aggregation, and up-sampling, among which, the selection of propagation and alignment strategies can significantly impact the performance[14, 15]. Therefore, here we concentrated on evaluating these two essential components, propagation and alignment. Specifically, we explored two mainstream propagation schemes, i.e., sliding window-based propagation and recurrent network-based propagation, and three representative alignment methods based on optical flow (OF), nonlocal attention (NA), and deformable convolution (DC). After validating these different schemes and methods (Fig. 1, Extended Data Figs. 2-4), we found that recurrent- and deformable convolution-based deep neural networks outperformed other combinations in aggregating cross-frame features and demonstrated greater potential for achieving superior super-resolution reconstruction for time-lapse biological data.

Although the recurrent network- and DC-based DNN architecture demonstrates convincing TISR performance, we noticed that the rudimentary designs in DC-based alignment limit the efficacy of information aggregation in two aspects. First, in live-cell fluorescence imaging, the acquisition frame rate is typically controlled carefully to a relatively low value to mitigate photobleaching or phototoxicity. Consequently, the motion-induced differences between two adjacent biological images are often larger than those observed in natural images. Deformable convolution is designed to learn additional offsets to allow the network to gather information beyond its regular local neighborhood[16]. Despite its offset diversity compared to conventional optical flow-based methods, DC still relies on relatively local convolutional operations and has a limited receptive field, producing unsatisfactory results when dealing with the large motion of biological structures. Second, DC-based alignment module is found to be difficult to train. The training instability of DC often results in offset overflow, deteriorating the final performance. To conquer this issue, Chan *et* al.[14] directly incorporated optical flow into the alignment module as initial offsets. However, in biological image TISR task, we adopted a similar method and observed that the optical flow learned by the network often fails to accurately present the actual motion of cell structures (Supplementary Fig. 1a). This discrepancy may be attributed to the heavier photon noise present in biological images compared to natural images, which substantially interferes with the accuracy of optical flow calculation and distorts the reconstruction results.

As delineated earlier, we contend that an effective alignment mechanism should possess several key attributes: (i) the capacity to model sub-pixel shifts and harness information from neighboring pixels and adjacent frames; (ii) the capability to accommodate substantial motion of biological structures, facilitating global alignment of neighboring frames; (iii) robustness against heavier photon noise, which can degrade optical flow estimation and compromise overall reconstruction quality.

To attain these objectives, especially for achieving global alignment and noise robustness, we direct our focus towards the Fourier domain. Firstly, every component within the Fourier frequency domain encapsulates information from the entire image, facilitating global analysis. Secondly, stochastic photon noise exhibits distinctive characteristics in the frequency domain, owing to its ultra-high frequency, enabling effective differentiation from the image signal. Our intuitive thinking is that alignment in frequency domain might be helpful for biological time-lapse image restoration.

In Fourier analysis, the translation of a signal can be expressed as the change of phase in the frequency domain while the amplitude remains unchanged, which is recognized as the frequency shifting property of Fourier transform. To illustrate this concept, let's consider a one-dimensional signal:

$$\mathrm{F}_t(\omega) = \mathrm{F}(\omega) * e^{-i\omega t} \tag{4}$$

where $\mathrm{F}(\omega)$ and $\mathrm{F}_t(\omega)$ represent the Fourier transform of the original signal and the signal after shifting by $t$ in spatial domain. Extending this concept into a two-dimensional image, the Fourier transform of a two-dimensional image after sptial-shifting can be denoted as:

$$\mathrm{F}_{x,y}(u,v) = \mathrm{F}(u,v) * e^{-i2\pi\left(\frac{ux}{\mathrm{M}}+\frac{vy}{\mathrm{N}}\right)}, \tag{5}$$

$$\left|\mathrm{F}_{x,y}(u,v)\right| = \left|\mathrm{F}(u,v)\right| \tag{6}$$

where $x, y$ represent two-dimensional translations and $\mathrm{M}, \mathrm{N}$ represent height and width of the image. It is evident that the translation of a signal is equivalent to adding a linear increment to its phase term in frequency space. In essence, by altering the phase of the Fourier transform of feature maps, we can reposition signals of different frequencies and achieve global movement of the feature maps. Notably, components of different frequencies can be relocated separately at a subpixel accuracy, enabling a diverse range of translation schemes.

Taken together, we designed the deformable phase-space alignment mechanism, which involves the following steps: (i) Apply Fourier transform on feature maps, i.e., current feature maps and neighbor feature maps to be aligned; (ii) Adaptively enhance the phase components via the phase convolution; (iii) Apply inverse Fourier transform with the enhanced phase and unchanged amplitude; (iv) Spatially warp the phase-enhanced feature maps using optical flow; (v) Generate offsets for deformable convolution; (vi) Apply deformable convolution to the feature maps for final alignment. See Fig. 2a and Extended Data Fig. 5 for the schematic illustration of DPA mechanism.

To test whether DPA outperforms conventional spatial DC mechanism and its potential variants, we conducted a comparative experiment involving four distinct alignment mechanisms, outlined as follows:

(I) Deformable phase-space alignment (DPA) and its variants

Real-valued fast Fourier transform (denoted as $\mathrm{FFT}(\cdot)$, implemented with *torch.fft.rfft*) is firstly applied on the current feature maps (denoted as $h_i$) and neighborhood feature maps (denoted as $h_{i-k}$):

$$h_i^{phase} = Angle(\mathrm{FFT}(h_i))$$

$$h_{i-k}^{phase} = Angle(\mathrm{FFT}(h_{i-k}))$$

$$h_i^{amplitude} = Abs\big(\mathrm{FFT}(h_i)\big)$$

$$h_{i-k}^{amplitude} = Abs\big(\mathrm{FFT}(h_{i-k})\big), \tag{7}$$

where $Angle(\cdot)$ and $Abs(\cdot)$ represent the operation to obtain element-wise angle and absolute value of the features.

(i) Amplitude convolution (Extended Data Fig. 6a): the concatenation of amplitudes, i.e., $h_i^{amplitude}$ and $h_{i-k}^{amplitude}$ goes through a convolutional layer (denoted as $f(\cdot)$) and two consecutive residual blocks (denoted as $\mathrm{RB}(\cdot)$) to learn the amplitude residual:

$$\delta(h_{i-k}^{amplitude}, h_i^{amplitude}) = \mathrm{RB}(\mathrm{RB}(f(p))) + h_{i-k}^{amplitude}$$

$$p = h_i^{amplitude} \oplus h_{i-k}^{amplitude}. \tag{8}$$

An inverse real-valued fast Fourier transform is utilized to reconstruct the amplitude-refined feature maps:

$$h_{i-k}^{refined,amplitude} = \mathrm{iFFT}\left(\delta(h_{i-k}^{amplitude}, h_{i}^{amplitude}) * e^{i*h_{i-k}^{phase}}\right). \quad (9)$$

(ii) Phase convolution (Extended Data Fig. 6b): the concatenation of phases, i.e., $h_i^{phase}$ and $h_{i-1}^{phase}$ goes through a convolutional layer and two consecutive residual blocks to learn the phase residual:

$$\delta(h_{i-k}^{phase}, h_{i}^{phase}) = \mathrm{RB}(\mathrm{RB}(f(p))) + h_{i-k}^{phase}$$

$$p = h_{i}^{phase} \oplus h_{i-k}^{phase}. \quad (10)$$

An inverse real-valued fast Fourier transform is utilized to reconstruct the phase-refine feature maps:

$$h_{i-k}^{refined,phase} = \mathrm{iFFT}\left(h_{i-k}^{amplitude} * e^{i*\delta(h_{i-k}^{phase}, h_{i}^{phase})}\right). \quad (11)$$

(iii) Phase & amplitude convolution (Extended Data Fig. 6c): both the concatenation of phase and the concatenation of amplitude go through a convolutional layer and a single residual block to learn the phase residual and amplitude residual, respectively.

$$\delta(h_{i-k}^{phase}, h_{i}^{phase}) = \mathrm{RB}(f(p_1)) + h_{i-k}^{phase}$$

$$p_1 = h_{i}^{phase} \oplus h_{i-k}^{phase}$$

$$\delta(h_{i-k}^{amplitude}, h_{i}^{amplitude}) = \mathrm{RB}(f(p_2)) + h_{i-k}^{amplitude}$$

$$p_2 = h_{i}^{amplitude} \oplus h_{i-k}^{amplitude}, \quad (12)$$

An inverse real-valued fast Fourier transform is utilized to reconstruct the phase & amplitude-refined feature maps:

$$h_{i-k}^{refined,phase\,\&\,amplitude} = \mathrm{iFFT}\left(\delta(h_{i-k}^{amplitude}, h_{i}^{amplitude}) * e^{i*\delta(h_{i-k}^{phase}, h_{i}^{phase})}\right). \quad (13)$$

(II) Deformable alignment (DA)

Spatial domain convolution (Extended Data Fig. 6d): the concatenation of current and neighborhood feature maps directly goes through a convolutional layer and two consecutive residual blocks to learn the spatial domain residual.

$$h_{i-k}^{refined,spatial-domain} = \mathrm{RB}(\mathrm{RB}(f(p))) + h_{i-k}$$

$$p = h_{i-k} \oplus h_{i}, \quad (14)$$

The refined feature maps are then fed into the deformable convolution module as is depicted in Extended Data Fig. 2 for subsequent spatial alignment.

We then examined four TISR models equipped with four alignment mechanisms detailed above on linear SIM data of MTs and simulated data of tubular structures with infallible GT references (Extended Data Fig. 6e-h). We found that feature alignment in the Fourier space typically outperforms alignment conducted solely in the spatial domain. Among the three frequential alignment mechanisms evaluated, phase convolution-based alignment consistently outperforms other configurations with comparable convolutional depth and computational complexity. This trend underscores the superiority of the proposed phase-space alignment mechanism.

#### 4. The design of Bayesian DPA-TISR

##### I. Reliability in deep neural networks

Within the domain of scientific investigation, the observation authenticity is of paramount importance. When the reliability of data cannot be confidently ascertained, it becomes evident that any subsequent conclusions drawn from such data are inherently susceptible to errors. This is particularly pertinent in the field of microscopy, where uncertainties in observations can lead to erroneous interpretations of biological phenomena.

This issue further exacerbates in the situation when computational techniques such as image restoration neural networks or the iterative deconvolution are adopted in the image processing pipeline. These advanced methods demonstrate exceptional efficacy in characterizing the statistical distributions of noises, resulting in notable enhancement in perceptual quality of the restored images. However, they also introduce a potential drawback: the risk of generating errors that closely resemble the ground truth, rendering them difficult to discern for scientists.

Therefore, in the development of deep learning-assisted microscopy technologies, as well as other fields, the evaluation and calibration of the confidence level of neural network outputs will play a significant role and have profound implications for future research and development. Particularly in the field of biological imaging, neural network models with quantifiable-confidence can assist biological researchers in better interpreting imaging results, thereby enhancing the rationality and accuracy of scientific conclusions drawn from the images.

Consequently, it is of vital for a computational image processing method to measure the uncertainty or confidence of restored images. Some efforts have been made in evaluating the dependability and uncertainty of neural networks, and find that the reliability of neural network predictions can be quantified through two predictive uncertainties: model uncertainty and data uncertainty akin to epistemic and aleatoric uncertainty, respectively, in Bayesian analysis[17-19]. Model uncertainty (epistemic uncertainty) encompasses the uncertainty arising from deficiencies in the model, including errors in the training process, inadequacies in the model's architecture, or the absence of knowledge due to unknown samples. Data uncertainty (aleatoric uncertainty), on the other hand, is associated with uncertainty that directly originates from the data itself. This type of uncertainty arises due to the loss of information when representing the real world within a data sample, including observation noise and imperfect ground truth. To mitigate model uncertainty, researchers typically collect more diversified training data or employ data augmentation techniques to expand the training set. Additionally, designing more robust network architectures and optimization algorithms can assist the network in better handling aleatoric uncertainty.

**II. Aleatoric uncertainty modelling**

In the task of deep learning-based image SR reconstruction, the acquisition of ground truth, i.e., the GT-SIM data in this work, is a crucial step. The GT-SIM images are typically obtained by reconstructing raw SIM images acquired using high laser power and long camera exposure. However, even when ensuring that biological samples do not move during acquisition, the inherent quantum nature of fluorescence and imperfect photoelectric conversion of the camera lead to different noise in each captured image and the corresponding ground truth image, contributing significantly to the aleatoric uncertainty in SR reconstructions of deep neural networks.

Inspired by previous work[20, 21], we assigned each pixel a Laplace distribution in the output SR image rather than a single intensity value:

$$p_{Laplace}(y;\hat{y},\hat{\sigma}) = \frac{1}{2\hat{\sigma}}\exp\left(-\frac{|y-\hat{y}|}{\hat{\sigma}}\right), \tag{15}$$

where $\hat{y}$ and $\hat{\sigma}$ denote the predicted value and scale of a certain pixel. In this way, the scale $\hat{\sigma}$ can be regarded as a measurement of the data uncertainty. Then the output SR image $\hat{y}$ and the scale $\hat{\sigma}$ can be simultaneously addressed by minimizing the negative log-likelihood (NLL) function:

$$\mathcal{L}_{NLL}(\hat{y},y) = \frac{1}{\mathrm{T}}\frac{1}{\mathrm{N}}\sum_{t=1}^{\mathrm{T}}\sum_{n=1}^{\mathrm{N}}\frac{|y_n^t-\hat{y}_n^t|}{\hat{\sigma}_n^t} + \log\hat{\sigma}_n^t \ , \tag{16}$$

where T denotes the number of timepoints and N denotes the number of pixels in a single image. Notably, with the scale parameters, the model has the flexibility to strike a balance between two extremes in minimizing the loss function above: achieving an accurate output prediction with low uncertainty or accommodating high uncertainty if an accurate output prediction is not feasible.

**III. Epistemic uncertainty modelling**

Compared to natural images, training data for microscopic images are more difficult to obtain. The diversity of biological specimens makes it challenging to collect a training dataset that encompasses all possible structures and fluorescent signal conditions. Furthermore, one of the primary objectives of biological imaging experiments is to discover new biological structures and bioprocesses, rendering the approach of mitigating cognitive uncertainty through extensive data collection impractical. In such cases, SR neural networks may not adequately learn all possible biological structures, leading to epistemic uncertainty.

As previously discussed, we address aleatoric uncertainty by predicting a probability distribution, as opposed to a single scalar value. However, this method cannot capture uncertainty associated with the model parameters $\theta$, which is induced by deficiencies in the model training procedure. For this purpose, we modified the DPA-TISR to a Bayesian neural network that employs a distribution over model parameters. Let $\{X, Y\}$ represent the training dataset. The posterior distribution over model parameters, denoted as $p(\theta|X, Y)$, can be expressed as follows:

$$p(\theta|X, Y) = \frac{p(Y|\theta, X)p(\theta)}{p(Y|X)} \tag{17}$$

Given a test sample $x^*$, the restored image $y^*$ can be predicted:

$$p(y^*|x^*, X, Y) = \int p(y^*|x^*, \theta)p(\theta|X, Y)d\theta \tag{18}$$

While this equation is mathematically complete, the computation of $p(\theta|X, Y)$cannot be carried out analytically but can be approximated by variational parameters $q(\theta)$. The objective is to approximate a distribution that closely resembles the posterior distribution obtained by the model. The degree of similarity between two distributions, measured by their KL divergence, can be expressed as follows:

$$KL(q(\theta)||p(\theta|X, Y)) = \int \log \frac{q(\theta)}{p(\theta|X, Y)} d\theta \tag{19}$$

KL divergence minimization can also be rearranged into the *evidence lower bound* (ELBO) maximization[22]:

$$L_{VI} := \int q(\theta) \log p(Y|X, \theta)\, d\theta - KL\big(q(\theta)\|p(\theta|X, Y)\big), \tag{20}$$

where the first term is dedicated to fitting the data, while the second term is geared towards approaching the prior distribution. This methodology is commonly referred to as variational inference (VI). Among various approaches, Dropout VI[20, 23] stands out as one of the most prevalent methods, which use dropout as a regularization term.

Leveraging the variational inference method, we can utilize a straightforward variational distribution $q(\theta)$ by incorporating concrete dropout[20] before each weight layer. This allows us to approximate the posterior distribution $p(\theta|X, Y)$. By employing Monte Carlo (MC) integration over M samples that adhere to $\theta^{(m)} \sim q(\theta)$, the equation above can be expressed as:

$$\mathrm{p}(y^*|x^*,\mathrm{X},\mathrm{Y}) \approx \int \mathrm{p}(y^*|x^*,\theta)\mathrm{q}(\theta)\mathrm{d}\theta \approx \frac{1}{\mathrm{M}}\sum_{\mathrm{m}=1}^{\mathrm{M}} \mathrm{p}\left(y^*\middle|x^*,\theta^{(\mathrm{m})}\right) \tag{21}$$

During the inference stage, the dropout layers in the Bayesian DPA-TISR randomly set input neurons to zero. By aggregating the outcomes of stochastic forward propagation through the trained model, the pixel-wise predictive mean can be computed as the super resolution result:

$$\hat{y}_{mean} = \frac{1}{\mathrm{M}}\sum_{m=1}^{\mathrm{M}} \hat{y}^{(m)}, \tag{22}$$

where $\hat{y}^{(m)}$ is the predicted mean from the $m^{th}$ network $\theta^{(\mathrm{m})}$. The model uncertainty is then quantified by calculated the standard deviation of the predicted results:

$$\sigma_{model} = \sqrt{\frac{1}{\mathrm{M}}\sum_{m=1}^{\mathrm{M}} (\hat{y}^{(m)} - \hat{y}_{mean})^2}\ . \tag{23}$$

Subsequently, the data uncertainty is assessed by computing the quadratic mean of the estimated variance:

$$\sigma_{data} = \sqrt{\frac{1}{\mathrm{M}}\sum_{m=1}^{\mathrm{M}} (\hat{\sigma}^{(m)})^2}\,, \tag{24}$$

where $\hat{\sigma}^{(m)}$ is the predicted scale map from the $m^{th}$ network $\theta^{(\mathrm{m})}$.

#### IV. Confidence calibration

Confidence is deemed as well-calibrated when its value accurately reflects the actual probability of correctness. Consequently, to leverage uncertainty quantification methods effectively, it is imperative to ensure that the network is well calibrated. For regression tasks, the calibration can be defined such that predicted confidence intervals should match the confidence intervals computed from the dataset, which can be effectively visualized through a reliability diagram. In practice, a DNN is considered under-confident when the predicted confidence is lower than the empirical accuracy (Supplementary Fig. SN1b), whereas it is regarded as over-confident when the predicted confidence exceeds the empirical accuracy (Supplementary Fig. SN1a). Well-calibrated metrics should yield credibility values similar to accuracy, resulting in a reliability diagram that is diagonal (Supplementary Fig. SN1c).

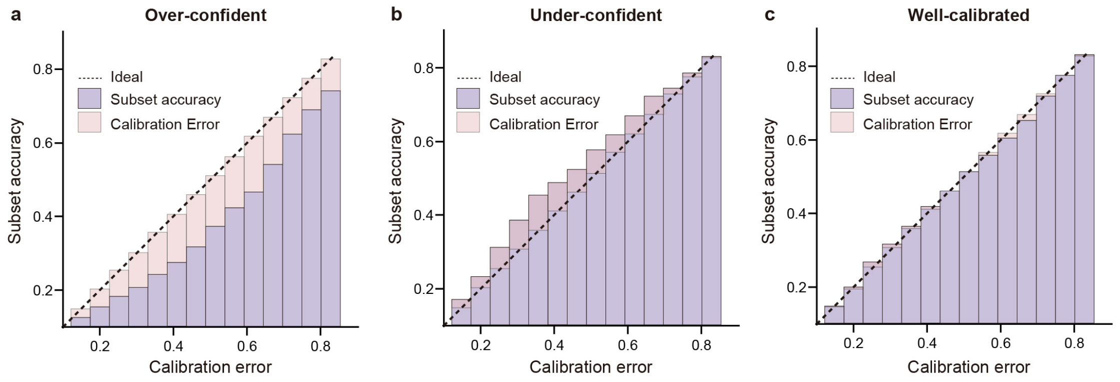

**Supplementary Figure SN1 | Typical reliability diagrams of DNN. a**, A representative reliability diagram of an over-confident model, where the subset accuracy is smaller than the corresponding confidence. **b**, A representative reliability diagram of an under-confident model, where the subset accuracy is larger than the corresponding confidence. **c**, A representative reliability diagram of a well-calibrated model, of which the subset accuracy is similar to the corresponding confidence.

Previous studies have highlighted that deeper networks often exhibit a tendency towards greater overconfidence compared to shallower ones[24]. After applying Bayesian neural networks to experimental data spanning multiple biological structures, we have similarly observed a propensity for estimated confidence levels to lean towards overconfidence (Extended Data Figs. 7-9). In addressing this confidence mismatch with the actual accuracy, various approaches have been employed, including regularization methods and post-processing techniques[24]. Post-processing methods typically necessitate a separate calibration dataset for calibration, and their effectiveness can be contingent upon the size of the validation dataset. Conversely, regularization methods entail modifications to the objective, optimization, and/or regularization procedures to develop inherently calibrated DNNs. In the context of biological image restoration tasks, our confidence correction method is structured upon regularization methods as is described in the Methods section of the main manuscript.

#### 5. Rolling Fourier ring correlation analysis

In conjunction with the confidence generated by Bayesian DPA-TISR deep neural network, the Fourier ring correlation (FRC)[25] was developed to evaluate the overall effective resolution of a fluorescence image by characterizing the highest reliable cut-off frequency in the Fourier domain. Based on FRC analysis, rolling Fourier ring correlation (rFRC)[26] was recently developed to provide the local distance measurements at the pixel level by rolling a sliding window across the image, which offers insights into the resolvability and uncertainty.

In our manuscript, we conducted the rFRC analysis as a comparative method in generating confidence map for TISR images. It was applied following the instructions of the original paper. First, an TISR image was down-sampled into two independent frames of identical contents with the same sampling rate by pixel shuffle, which is commonly used operation to augment the raw data in FRC analysis. Next, the input images were padded symmetrically around a half size of the block to ensure accurate FRC calculation at the image boundaries. Then a background threshold was set for the center pixel to avoid FRC calculation in background areas. If the mean of the center pixels exceeded the threshold, the FRC was computed within a 32-pixel size block (which is the default size of the software), and the resulting FRC resolution was assigned to the center pixel of each block. Conversely, if the mean was below the threshold, a zero value was assigned to the central pixel. This process was repeated block by block across the entire image to generate a final rFRC confidence map.

### Supplementary Figures

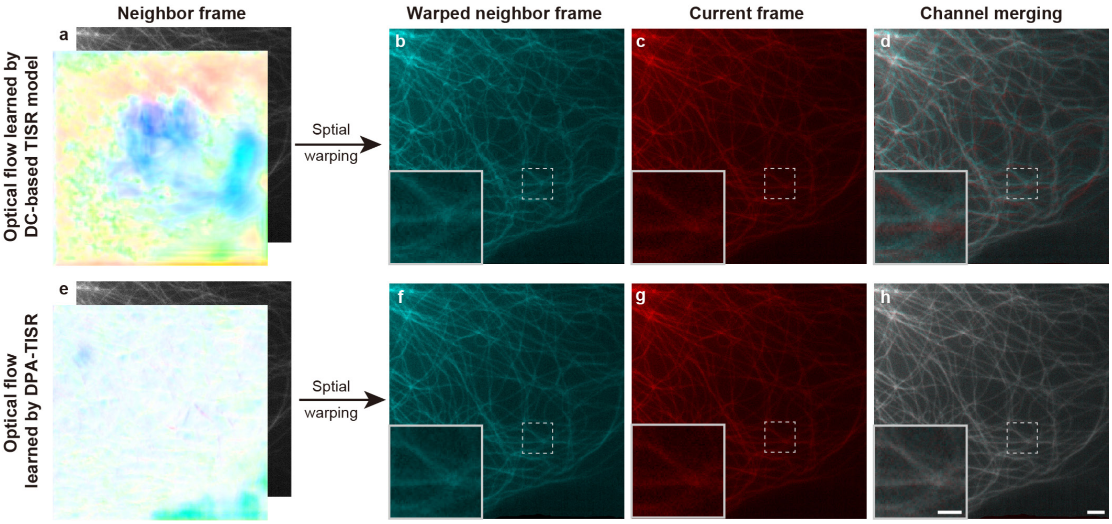

**Supplementary Fig. 1 | Visualization of optical flow-based spatial warping.** **a**, Wide-field neighbor frame of microtubules and corresponding optical flow relative to the current frame generated via a deformable convolution-based TISR model. **b**, The warped neighbor frame using the optical flow shown in a. **c**,**d**, Channel merging visualization (d) of the warped neighbor frame (b) and current frame (c). **e**, Wide-field neighbor frame of microtubules and corresponding optical flow relative to the current frame generated via the DPA-TISR model. **f**, The warped neighbor frame using the optical flow shown in e. **g**,**h**, Channel merging visualization (h) of the warped neighbor frame (f) and current frame (g). These results illustrate that the phase-space alignment mechanism in the DPA-TISR model can synergize with optical flow calculation, resulting in more accurate and finer feature alignment. Scale bar: 3 µm, 1 µm (zoom-in regions).

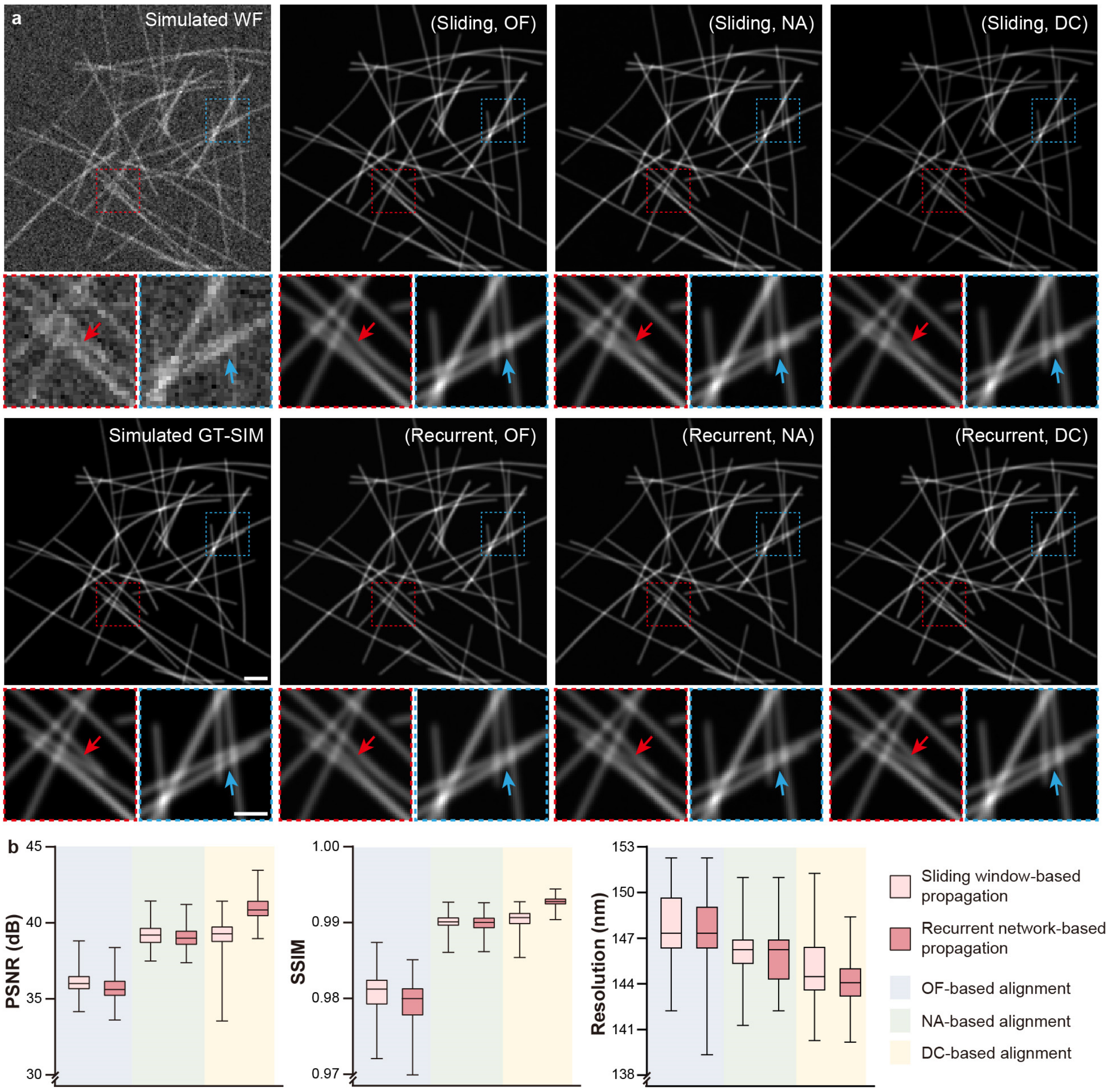

**Supplementary Fig. 2 | Comparison of representative propagation and alignment mechanisms in TISR models with simulated data of high SNR ($\alpha = 20$) and low framerate (3 Hz). a**, Representative TISR images of simulated tubular structures inferred by six models combined by two propagation methods, sliding window-based propagation and recurrent network-based propagation, and three alignment mechanisms based on optical flow (OF), nonlocal attention (NA) and deformable convolution (DC). WF and GT-SIM images are shown in the first column for reference. **b**, Statistical comparison of the six models in terms of PSNR, SSIM, and resolution quantified by decorrelation analysis[27] (n=200). Center line, medians; limits, 75% and 25%; whiskers, maximum and minimum. Scale bar, 1 μm (a), 0.5 μm (zoom-in regions in a).

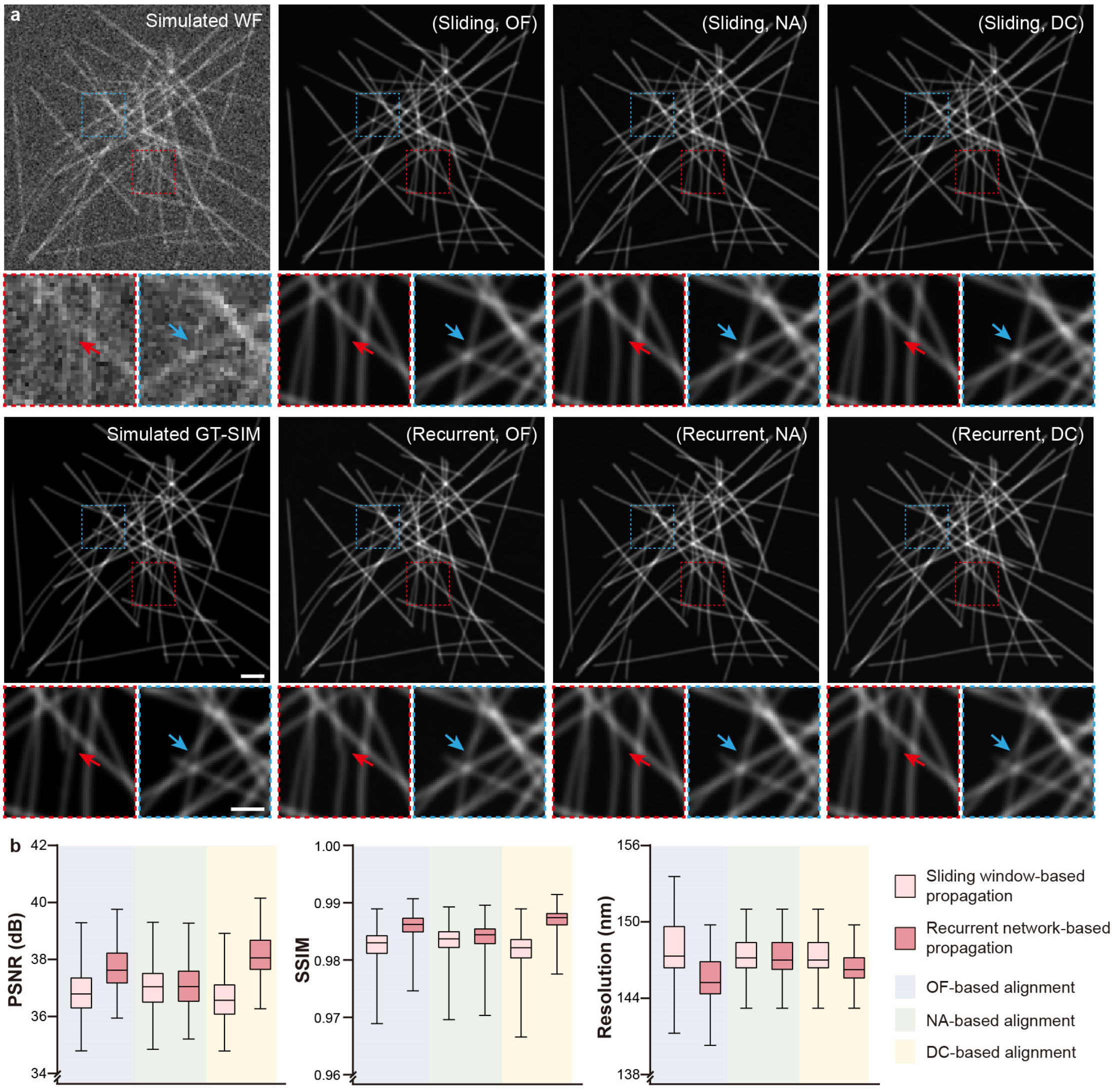

**Supplementary Fig. 3 | Comparison of representative propagation and alignment mechanisms in TISR models with simulated data of high SNR ($\alpha = 20$) and high framerate (30 Hz). a**, Representative TISR images of simulated tubular structures inferred by six models combined by two propagation methods, sliding window-based propagation and recurrent network-based propagation, and three alignment mechanisms based on optical flow (OF), nonlocal attention (NA) and deformable convolution (DC). WF and GT-SIM images are shown in the first column for reference. **b**, Statistical comparison of the six models in terms of PSNR, SSIM, and resolution quantified by decorrelation analysis[27] (n=200). Center line, medians; limits, 75% and 25%; whiskers, maximum and minimum. Scale bar, 1 μm (a), 0.5 μm (zoom-in regions in a).

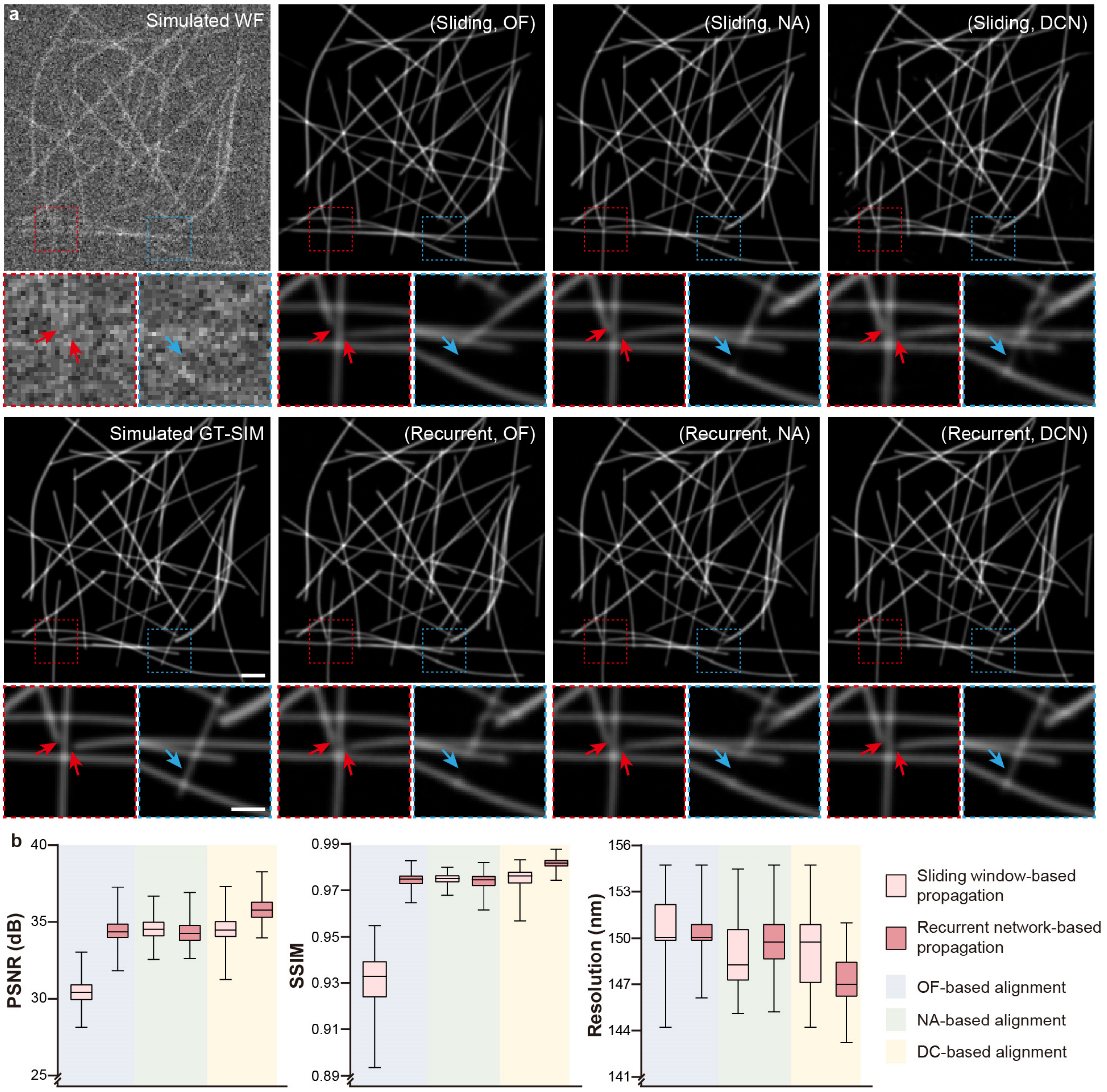

**Supplementary Fig. 4 | Comparison of representative propagation and alignment mechanisms in TISR models with simulated data of low SNR ($\alpha = 5$) and low framerate (3 Hz). a**, Representative TISR images of simulated tubular structures inferred by six models combined by two propagation methods, sliding window-based propagation and recurrent network-based propagation, and three alignment mechanisms based on optical flow (OF), nonlocal attention (NA) and deformable convolution (DC). WF and GT-SIM images are shown in the first column for reference. **b**, Statistical comparison of the six models in terms of PSNR, SSIM, and resolution quantified by decorrelation analysis[27] (n=200). Center line, medians; limits, 75% and 25%; whiskers, maximum and minimum. Scale bar, 1 µm (a), 0.5 µm (zoom-in regions in a).

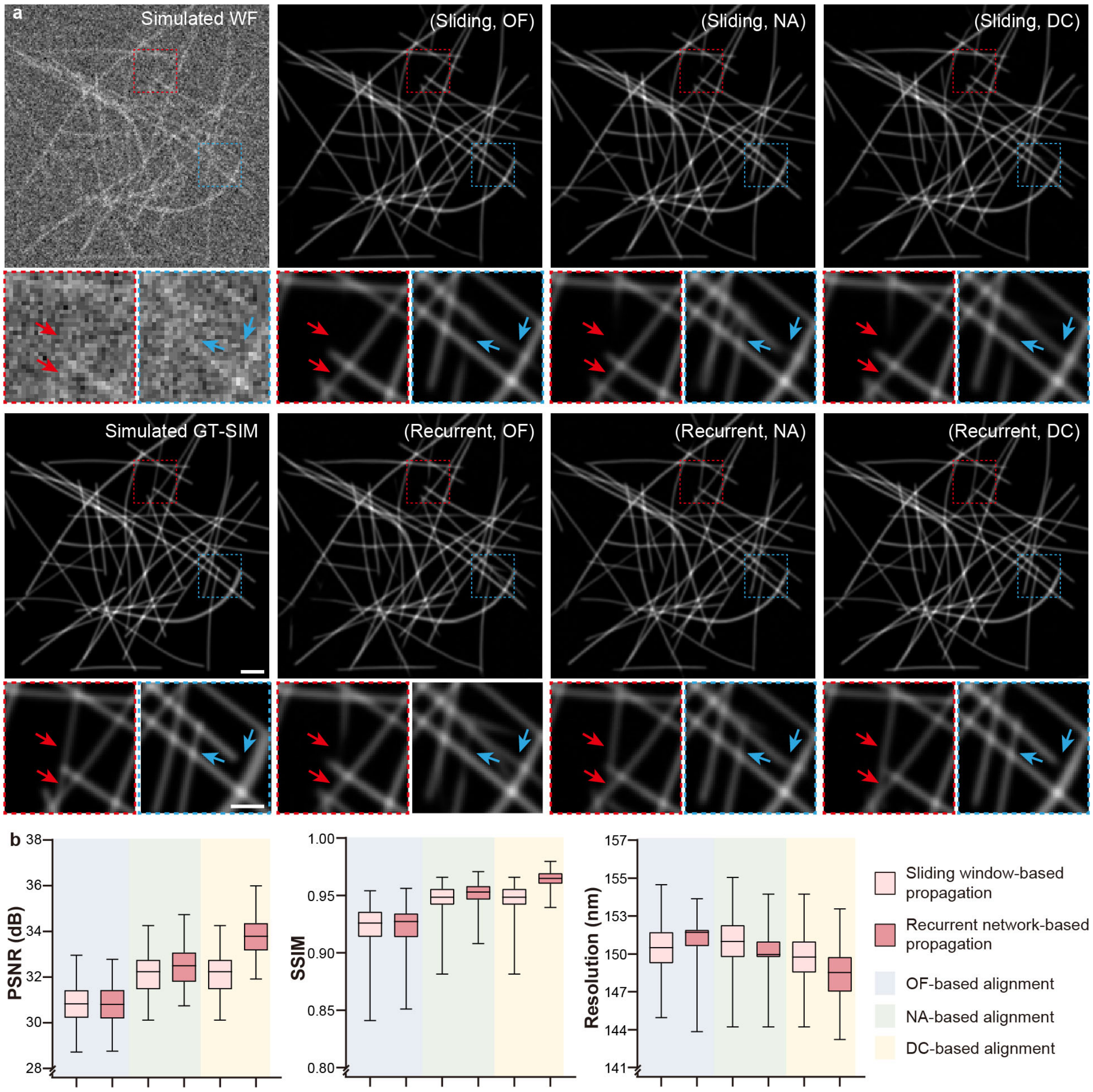

**Supplementary Fig. 5 | Comparison of representative propagation and alignment mechanisms in TISR models with simulated data of low SNR ($\alpha = 5$) and high framerate (30 Hz). a**, Representative TISR images of simulated tubular structures inferred by six models combined by two propagation methods, sliding window-based propagation and recurrent network-based propagation, and three alignment mechanisms based on optical flow (OF), nonlocal attention (NA) and deformable convolution (DC). WF and GT-SIM images are shown in the first column for reference. **b**, Statistical comparison of the six models in terms of PSNR, SSIM, and resolution quantified by decorrelation analysis[27] (n=200). Center line, medians; limits, 75% and 25%; whiskers, maximum and minimum. Scale bar, 1 µm (a), 0.5 µm (zoom-in regions in a).

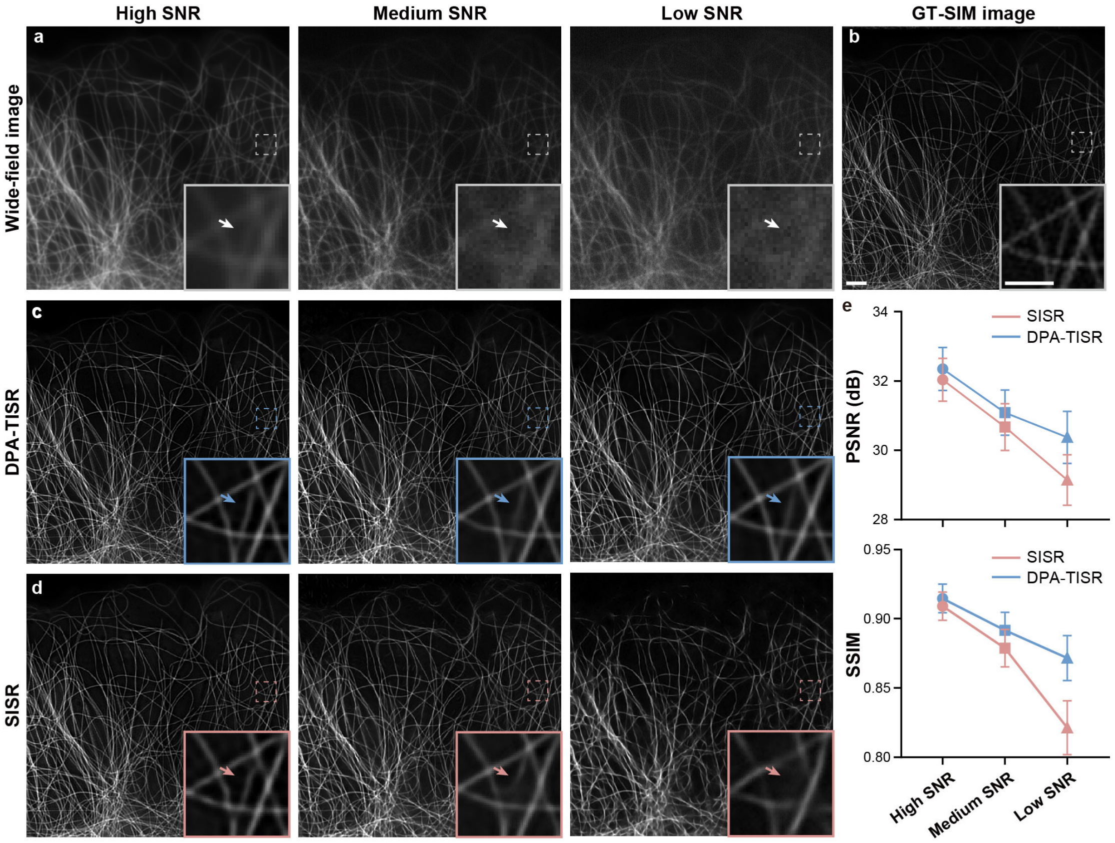

**Supplementary Fig. 6 | Comparison of DPA-TISR and SISR models with microtubule images under different SNR conditions. a**, Wide-field images of high (first column), medium (second column) and low SNR (third column) from the BioTISR dataset. **b**, GT-SIM image of the same region shown in a. **c**,**d**, SR results inferred from WF images via DPA-TISR (c) and its SISR (d) variant (see Methods section in the main manuscript for details of the modification). **e,** Statistical comparison of DPA-TISR and SISR models in terms of PSNR (upper panel) and SSIM (lower panel) calculated at 3 levels of SNR for microtubule data (n=50). Central point, medians; whiskers, mean±standard deviation. Scale bar: 3 μm, 1 μm (zoom-in regions).

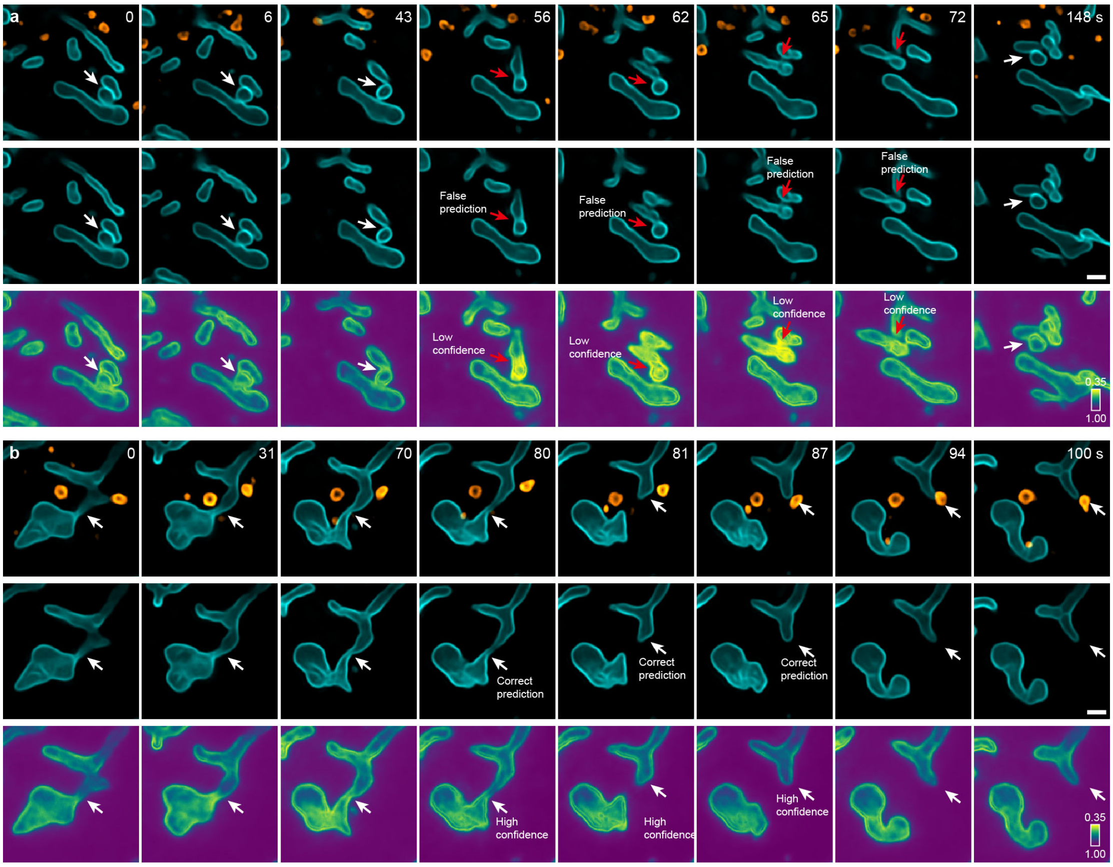

**Supplementary Fig. 7 | Representative correct and false predictions indicated by the confidence map generated via Bayesian DPA-TISR. a**, Time-lapse SR images of mitochondrial and lysosomes, as well as corresponding confidence map of mitochondrial images generated by Bayesian DPA-TISR, showcasing an equivocal mitochondrial fusion event. The red arrows indicate the potential false predictions of DPA-TISR identified by the low confidence value, which alerts us it may not be a real fusion event. **b**, Time-lapse SR images of mitochondrial and lysosomes, as well as corresponding confidence map of mitochondrial images generated by Bayesian DPA-TISR, showcasing a typical mitochondrial fission event. The high confidence of the prediction indicates a high reliability of this observation. Scale bar, 1 μm (a, b).

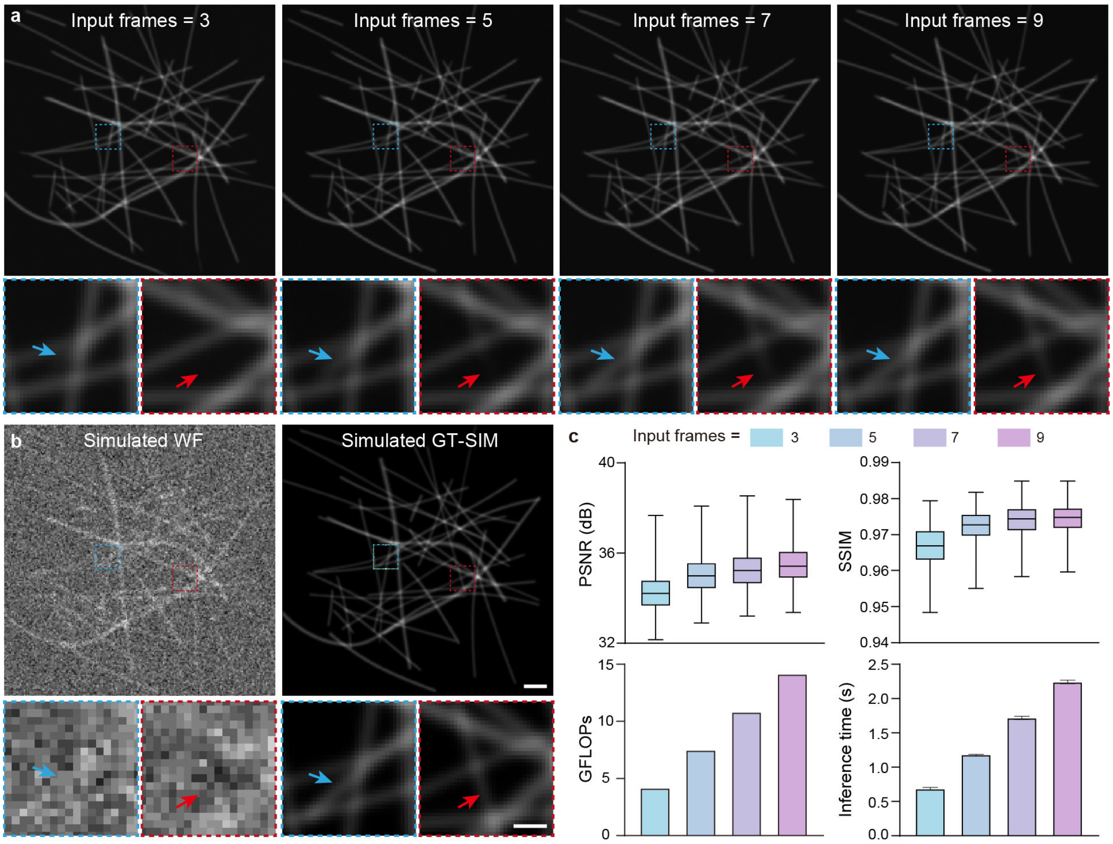

**Supplementary Fig. 8 | Evaluation of DPA-TISR trained with different input configuration. a**, TISR images inferred by DPA-TISR models trained and tested with different input sequence length ranging from 3 to 9. **b**, Corresponding simulated wide-field image as inputs and GT-SIM image shown for reference. **c**, Statistical comparison of DPA-TISR models trained and tested with different input sequence length in terms of PSNR, SSIM, GFLOPs and inference time on simulated data (n=200). For the box plots in the upper row: center line, medians; limits, 75% and 25%; whiskers, maximum and minimum. These evaluations demonstrate that the performance of DPA-TISR is generally enhanced with increasement of the input sequence length, however, at the expense of higher computation complexity. According to the evaluation, we selected an input sequence length of 7 in our experiments of this paper for compromise between efficiency and performance. Scale bar, 1 μm, 0.25 μm (zoom-in regions).

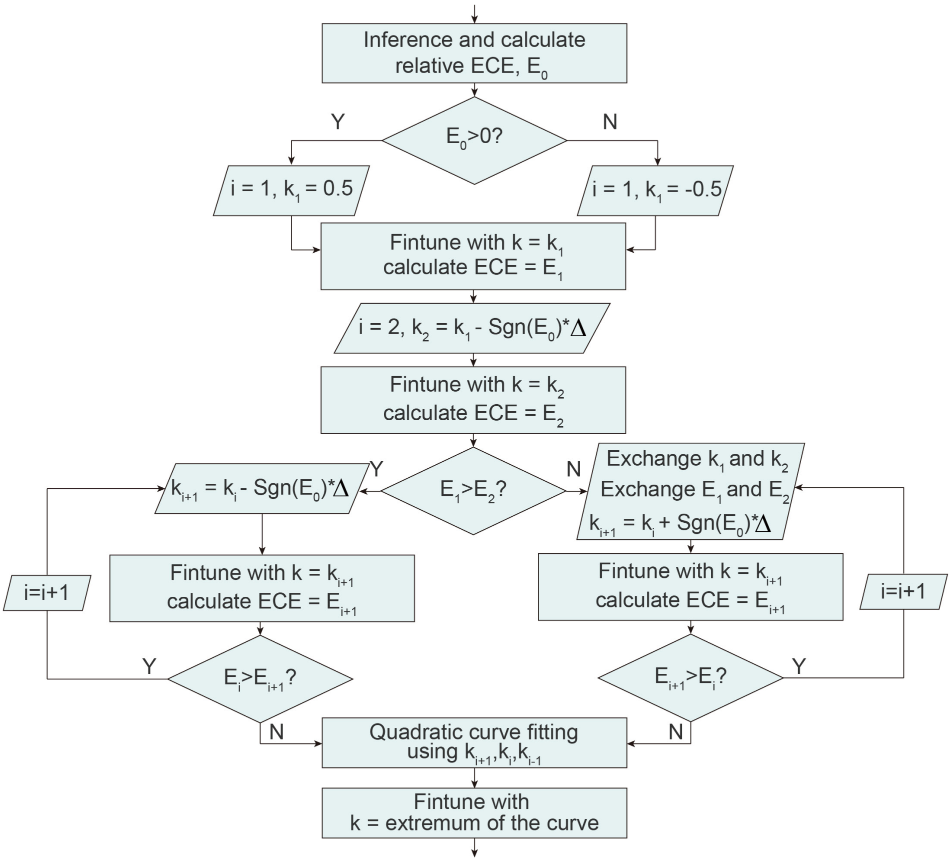

**Supplementary Fig. 9 | Flow diagram of the confidence calibration procedure.**

#### Captions for Supplementary Videos

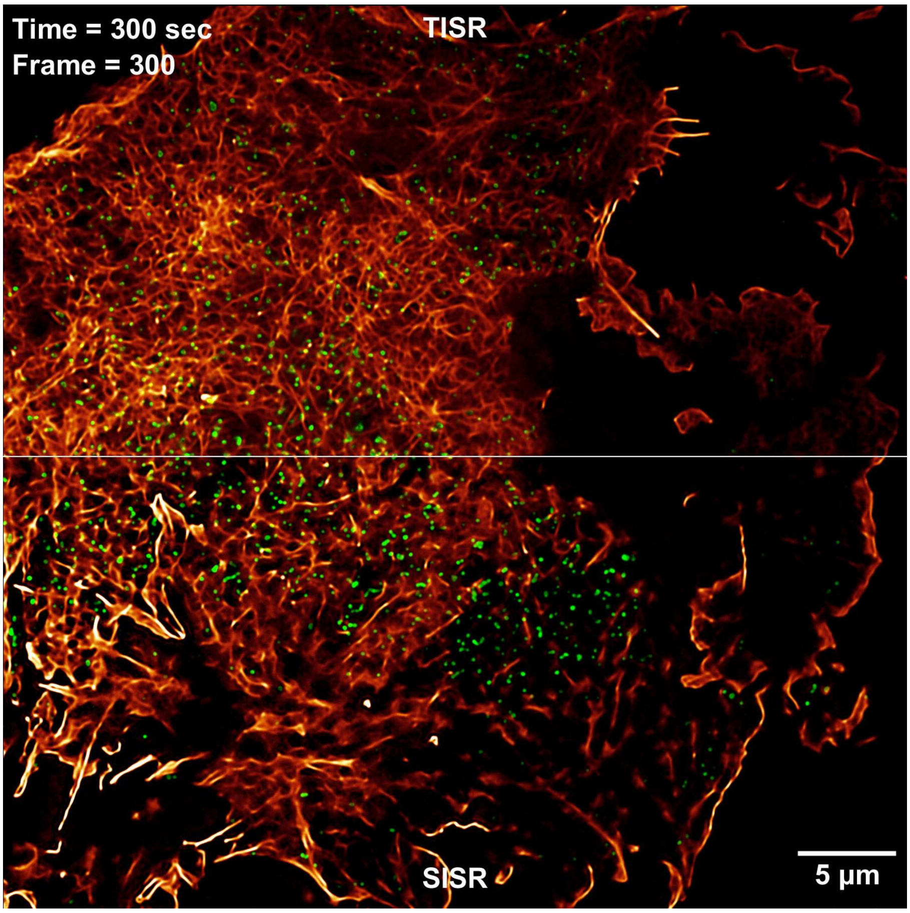

**Supplementary Video 1 | Two-color SR live-imaging of CCPs and F-actin enabled by DPA-TISR.** DPA-TISR enhanced TIRF imaging of CCPs and F-actin over 4,800 timepoints at 1-sec interval of a COS-7 cell co-expressing Lifeact-mEmerald (red) and Clathrin-mCherry (green), visualizing the rapid dynamics and interactions between the two subcellular structures.

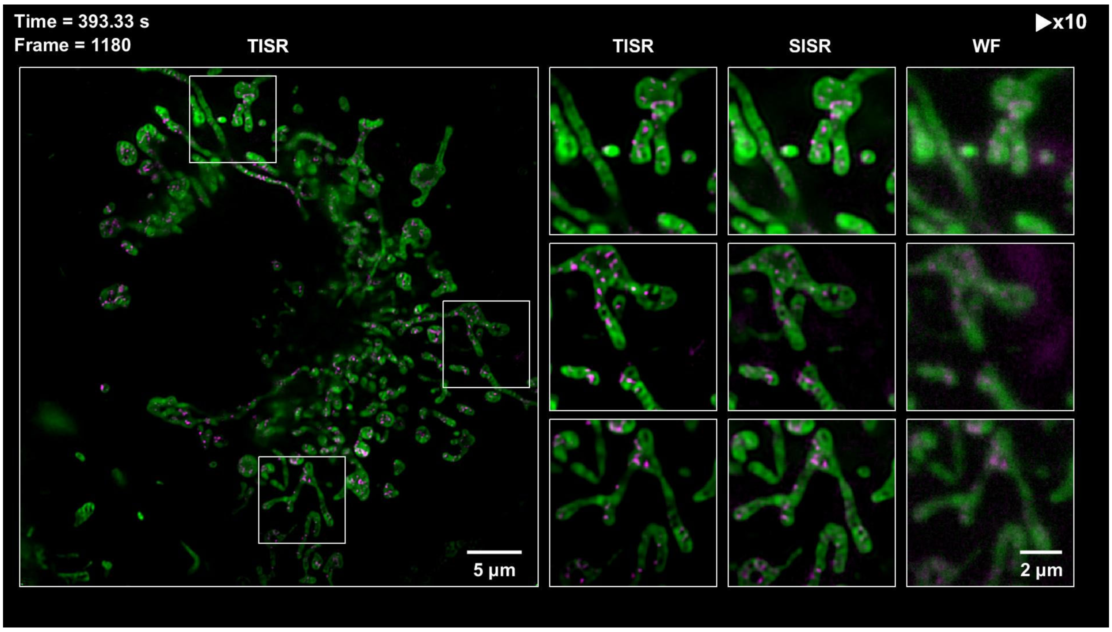

**Supplementary Video 2 | Long-term SR recordings of mitochondrial cristae and nucleoid dynamics with DPA-TISR.** Two-color SR live-imaging of a COS-7 cell labelled with PKMO-Halo (green) and TFAM-mEmerald (magenta) by DPA-TISR over 1,680 timepoints at a framerate of 3Hz, recoding mitochondrial fission and fusion and reorganizations of nucleoids concomitant with cristae deformation.

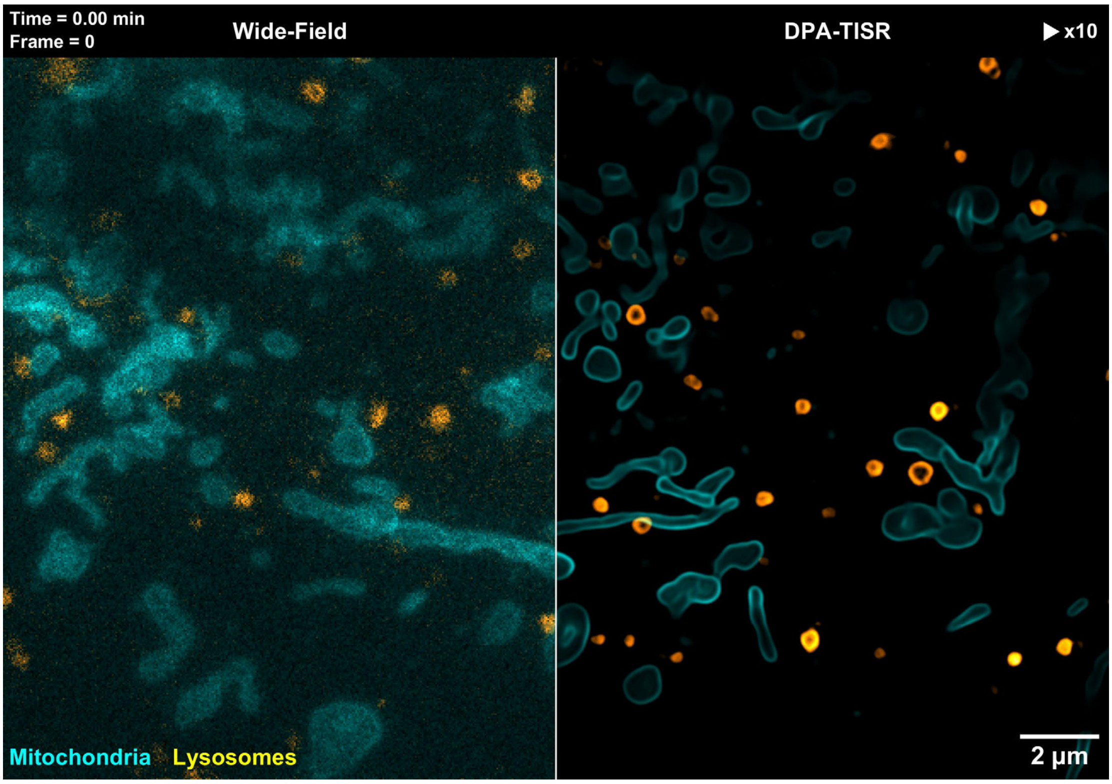

**Supplementary Video 3 | Long-term SR live-imaging of outer mitochondrial membrane and lysosomes enabled by DPA-TISR.** Two-color SR live-imaging of a COS-7 cell labelled with 2×mEmerald-Tomm20 (cyan) and Lamp1-Halo (yellow) over a ultralong time course of ~10,000 timepoints at 0.5-second intervals, showcasing multiple lysosome-mediated mitochondrial dynamics such as hitchhiking and fissions.

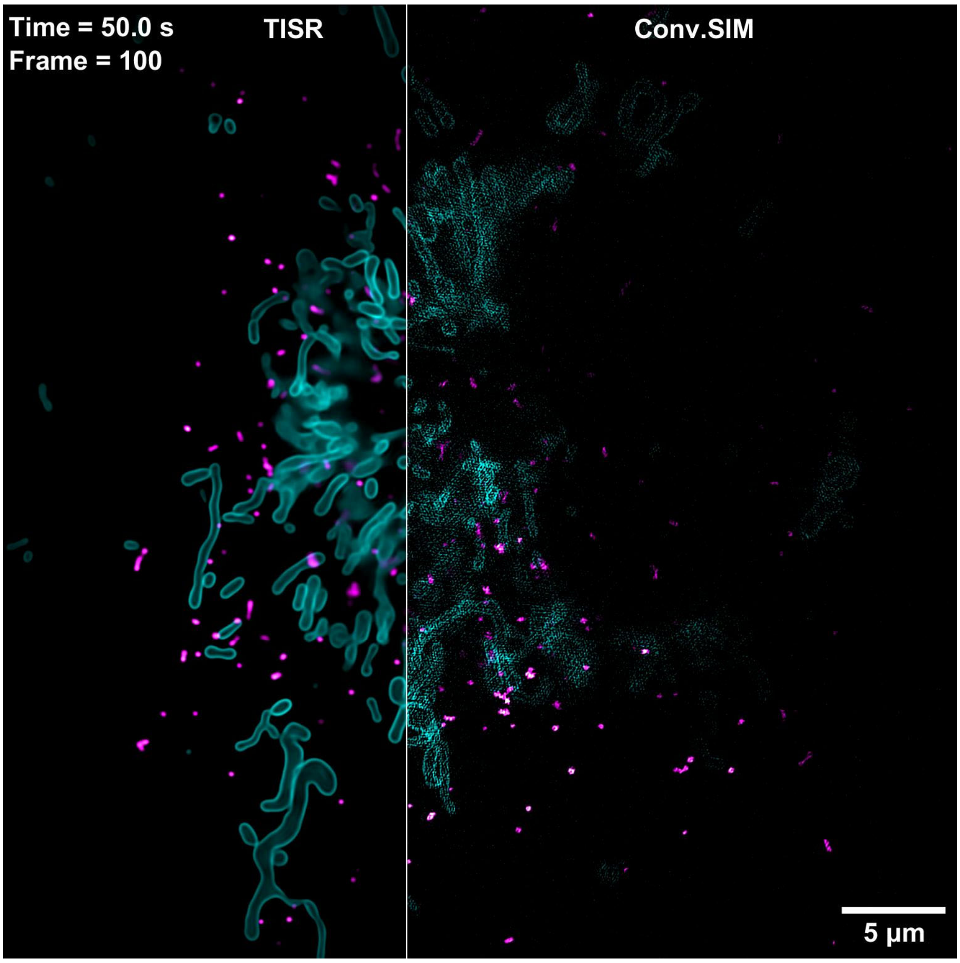

**Supplementary Video 4 | Dynamic interactions between mitochondria (Mito) and peroxisomes (PO) visualized by DPA-TISR.** SR live-imaging reconstructed via DPA-TISR of a COS-7 cell co-expressing 2×mEmerald-Tomm20 (cyan) and PMP-Halo (magenta) with high spatiotemporal resolution and long duration, revealing intricacy of complex Mito-PO contact behaviors.

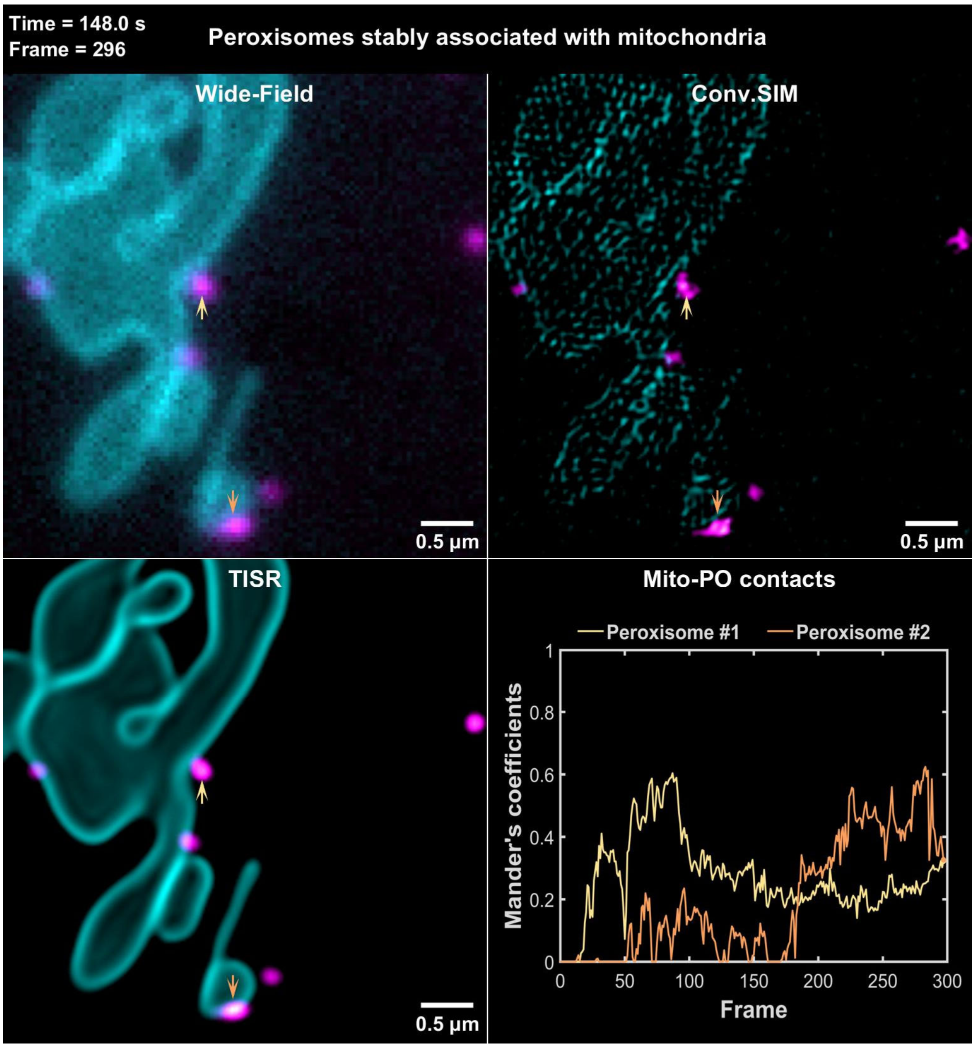

**Supplementary Video 5 | Examples of different types of Mito-PO contacts.** Representative examples of four types of Mito-PO contacts: (i) POs stably associated with a single Mito; (ii) POs served as the bridge that simultaneously tethered two Mito; (iii) POs unexpectedly changing their contact sites from one Mito to another; and (iv) POs with no contact with Mito. Wide-field images (top left corner), conventional SIM images (top right corner), DPA-TISR images reconstructed from WF images (bottom left corner) are displayed for comparison. The Mander's overlap coefficient plots are shown in the bottom right panel, which serves as a quantitative indicator of the strength of Mito-PO contacts.
